# Arrestin-3-assisted activation of JNK3 mediates dopaminergic behavioral and signaling plasticity in vivo

**DOI:** 10.1101/2023.10.27.564447

**Authors:** Mohamed R. Ahmed, Chen Zheng, Jeffery L. Dunning, Mohamed S. Ahmed, Connie Ge, F. Sanders Pair, Vsevolod V. Gurevich, Eugenia V. Gurevich

## Abstract

In rodents with unilateral ablation of the substantia nigra neurons supplying dopamine to the striatum, chronic treatment with the dopamine precursor L-DOPA or dopamine agonists induces a progressive increase of behavioral responses, a process known as behavioral sensitization. The sensitization is blunted in arrestin-3 knockout mice. Using virus-mediated gene delivery to the dopamine-depleted striatum of arrestin-3 knockout mice, we found that the restoration of arrestin-3 fully rescued behavioral sensitization, whereas its mutant defective in JNK activation did not. A 25-residue arrestin-3-derived peptide that facilitates JNK3 activation in cells, expressed ubiquitously or selectively in the direct pathway striatal neurons, fully rescued sensitization, whereas an inactive homologous arrestin-2-derived peptide did not. Behavioral rescue was accompanied by the restoration of JNK3 activity and of JNK-dependent phosphorylation of the transcription factor c-Jun in the dopamine-depleted striatum. Thus, arrestin-3-dependent JNK3 activation in direct pathway neurons is a critical element of the molecular mechanism underlying sensitization.

## Introduction

Signaling via G protein-coupled receptors (GPCRs) is controlled by a conserved two-step homologous desensitization mechanism: phosphorylation of the active receptors by GPCR kinases and subsequent binding of arrestins, which preclude further G protein activation (Gurevich and Gurevich, 2006). Two non-visual arrestin isoforms, arrestin-2 and arrestin-3 (Arr3) (a.k.a. β-arrestin1 and β-arrestin2, respectively), are ubiquitously expressed and negatively regulate the signaling of numerous GPCRs. Arrestins also act as positive regulators of cellular signaling via the assembly of multi-protein complexes (Gurevich and Gurevich, 2019, 2023; Peterson and Luttrell, 2017; Wess et al., 2023). The best-known signaling activity of arrestins is their ability to facilitate the activation of the mitogen-activated protein kinases ERK and JNK (Peterson and Luttrell, 2017; Wess et al., 2023). Reduced availability of arrestins or their complete deletion in cultured cells or living animals results in augmented G protein-mediated responses, reflecting defective GPCR desensitization (Bohn et al., 2003; Bohn et al., 1999; Gainetdinov et al., 1999). However, mice lacking arrestins also manifest loss-of-function phenotypes, suggesting the involvement of arrestin-mediated signaling. Mice lacking arrestins demonstrate blunted behavioral responses to dopaminergic drugs (Beaulieu et al., 2005; Urs et al., 2016; Zurkovsky et al., 2017), which suggests a role for arrestin-dependent signaling in the dopaminergic control of behavior.

Chronic administration of many drugs targeting GPCRs causes long-lasting adaptations. The best-known adaptation is tolerance, when the response to the drug diminishes with repeated administration (Colvin et al., 2019; Li, 2016; Qiao et al., 2012). Some drugs also induce the opposite adaptation, referred to as sensitization or reversed tolerance, i.e., enhanced response with repeated use (Cenci, 2007; Li, 2016; Qiao et al., 2012; Sgambato-Faure et al., 2005), or cause a paradoxical enhancement of the initial symptoms, as in opioid-induced hyperalgesia (Colvin et al., 2019). Molecular mechanisms that might underlie tolerance operate in cultured cells, while sensitization in any form is only observed in living animals. Both types of adaptations often coexist *in vivo*, with some drug effects displaying tolerance and others sensitization. Long-term adaptations, or mis-adaptations, during drug treatment present a serious clinical problem limiting the efficacy of the therapy and/or causing detrimental side effects.

The brain dopaminergic system is critical for the control of motor behavior, reward mechanisms, and cognition. The highest density of the dopaminergic innervation and the highest concentration of dopamine (DA) receptors are found in the striatum, a component of the subcortical complex of structures, which plays an essential role in movement control, motivation, and reward. All five DA receptor subtypes are GPCRs and interact with arrestins, although the mode of interaction and functional consequences are unique for each subtype (Burström et al., 2023; Kim, 2023; Kim et al., 2001; Li et al., 2015; Moritz et al., 2023; Yang et al., 2022). Long-term use of dopaminergic drugs causes persistent changes in dopamine-dependent behaviors. In rodents with unilateral ablation of the dopaminergic input to the striatum, stimulation with dopaminergic agonists or the DA precursor L-DOPA causes persistent behavioral and molecular sensitization [reviewed in (Bastide et al., 2015)]. The behavioral sensitization can be elicited in intact animals upon chronic treatment by diverse dopaminergic drugs such as psychostimulants and antipsychotics (Delage et al., 2023; Jeffery and Peter, 2011; Li, 2016; Qiao et al., 2012). The molecular mechanisms of long-term behavioral and signaling plasticity associated with persistent dopaminergic stimulation remain elusive.

Here we show that Arr3 is indispensable for the behavioral sensitization to L-DOPA in mice with the unilateral dopaminergic depletion and that its action is mediated by the Arr3-dependent activation of JNK in the striatal direct pathway medium spiny neurons (MSNs). This is an excellent mouse model of a long-term dopaminergic behavioral plasticity suitable for mechanistic studies of signaling. It is also a widely used animal model of Parkinson’s disease (PD). Behavioral alterations caused by L-DOPA in these mice bear an uncanny resemblance to L-DOPA-induced dyskinesia (LID), a side-effect of the dopamine replacement therapy in PD, which makes it an animal model of LID. Our findings reveal the role of Arr3-dependent JNK activation in the molecular mechanism of sensitization, thereby identifying it as a novel target for anti-LID therapy.

## RESULTS

### The loss of Arr3 suppresses behavioral sensitization to L-DOPA

Rodents with unilateral 6-hydroxydopamine (6-OHDA) lesions of substantia nigra neurons lack the dopaminergic innervation of the striatum ipsilateral to the lesion and respond by contralateral rotations to the DA precursor L-DOPA or dopaminergic agonists (Bastide et al., 2015). Chronic treatment of these animals with L-DOPA causes behavioral sensitization, i.e., progressively increased frequency of rotations. We examined sensitization of the L-DOPA-induced rotational behavior in 6-OHDA-lesioned arrestin-3 (A3KO) and arrestin-2 (A2KO) knockout mice as compared to wild type (WT) littermates. We found that A2KO mice showed an initial tendency to reduced rotation frequency, but eventually reached the same level as WT (**Fig. 1A**). In contrast, A3KO mice demonstrated reduced rotation frequency and minimal sensitization to L-DOPA (**Fig. 1A**). A3KO mice were previously shown to have reduced behavioral sensitivity (Beaulieu et al., 2005; Zurkovsky et al., 2017) and locomotor sensitization (Zurkovsky et al., 2017) to amphetamine. Therefore, their blunted response to L-DOPA could reflect an overall lower sensitivity to dopaminergic agents. To test whether this was the case, we compared locomotor responses to a one-time administration of the DA agonist apomorphine in WT and A3KO mice and found them to be similar (**Fig. 1B**). Next, we performed the cylinder test. Loss of DA upon 6-OHDA lesion reduces the use of the contralateral paw affected by the lesion to support the body, whereas L-DOPA supplies DA to the DA-depleted striatum and increases its use. We found that L-DOPA improved the use of the affected paw to a similar extent in WT and A3KO mice, indicating that this acute response to L-DOPA was preserved in A3KO animals (**Fig. 1C**). Thus, generally reduced dopaminergic signaling does not underlie the sensitization defect of A3KO mice.

**Figure 1.**
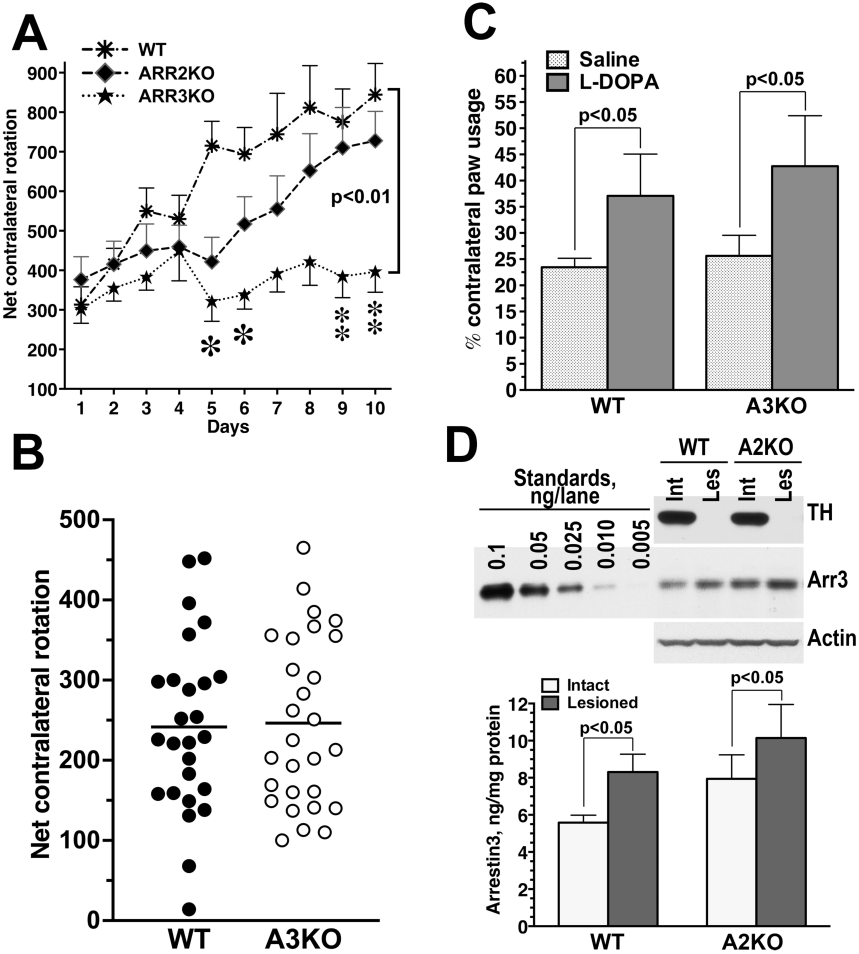
Hemiparkinsonian mice lacking Arr3 (A3KO) respond to dopaminergic stimulation but do not develop behavioral sensitization to L-DOPA. **(A)** Arrestin-2 knockout (A2KO), A3KO, and WT littermates with unilateral 6-hydroxydopamine lesions were chronically treated with L-DOPA and tested for L-DOPA-induced rotations. Means±S.E.M. are shown. The data were analyzed by two-way repeated measure ANOVA with GENOTYPE as between group and DAY as within group factor. The statistical comparison was made with respective littermates. WT littermates of A2KO and A3KO were similar. A2KO mice showed slightly reduced rotations (not statistically significant). A3KO mice demonstrated significantly reduced rotation frequency (Genotype effect F(1,162)=11.7, p=0.003, GENOTYPE X DAYF(9,162)=3.246, p=0.0012), indicative of impaired behavioral sensitization to L-DOPA. * -p<0.05, ** -p<0.01 by post hoc unpaired Student’s t-test for individual days. **(B)** Apomorphine-induced contralateral rotations in WT and A3KO mice. No difference was detected. **(C)** Hemiparkinsonian A3KO and WT mice were tested for forelimb usage in the cylinder test. The mice were injected with saline or L-DOPA (5 mg/kg s.c.) on separate days in a counterbalanced manner. The percentage of contralateral paw usage out of the total is shown. The data were analyzed by two-way repeated measure ANOVA with GENOTYPE as between group and L-DOPA as within group factor. The effect of L-DOPA was significant (p=0.032) whereas the effect of GENOTYPE was not (p=0.89). **(D)** <u>Upper panel</u>: Representative Western blot showing the expression of arrestin-3 in the intact and lesioned striatum in WT and A2KO mice. <u>Lower panel:</u> Quantification of the Western blot data demonstrating a significant upregulation of Arr3 in the lesioned hemisphere in both genotypes (Hemisphere within group factor across genotypes p=0.0005). There was no significant difference between WT and A2KO (p=0.2).

We previously detected an upregulation of Arr3 in the affected hemisphere in 6-OHDA-lesioned rats upon chronic L-DOPA treatment (Ahmed et al., 2007). We confirmed this finding in WT mice and found a similar increase in A2KO mice (**Fig. 1D**). Although there was a tendency towards upregulation of Arr3 in both hemispheres of A2KO mice as compared to WT, it did not reach statistical significance (**Fig. 1D**). Collectively, these data suggest that Arr3 is required for the sensitization, and its elevated expression upon L-DOPA treatment might contribute to the development of locomotor sensitization to L-DOPA.

### Exogenous Arr3 in the striatum rescues behavioral sensitization in A3KO mice

When unilaterally lesioned rodents are treated with a high dose of L-DOPA, they display contralateral rotations, which increase in frequency with each drug administration (Ahmed et al., 2010; Ahmed et al., 2007; Bastide et al., 2015; Bychkov et al., 2007). They also show other types of induced movements collectively referred to as **<u>A</u>**bnormal **I**nvoluntary **<u>M</u>**ovements (AIMs) (Ahmed et al., 2010; Cenci and Lundblad, 2007), the frequency of which progressively increases, demonstrating sensitization. The loss of Arr3 reduces sensitization measured by rotations (**Figs. 1A**, **2A**) and AIMs (**Fig. 2B**), suggesting that Arr3 is an important contributor to behavioral sensitization. Germline deletion of Arr3 could have resulted in developmental alterations in the brain circuitry and/or signaling mechanisms that ultimately affect the brain plasticity. To test whether Arr3 acts during the development of sensitization, we employed a rescue strategy with lentivirus-mediated gene transfer. Lentivirus (LV) encoding HA-tagged WT Arr3 was used to restore Arr3 in the DA-depleted striatum of A3KO mice.

**Figure 2.**
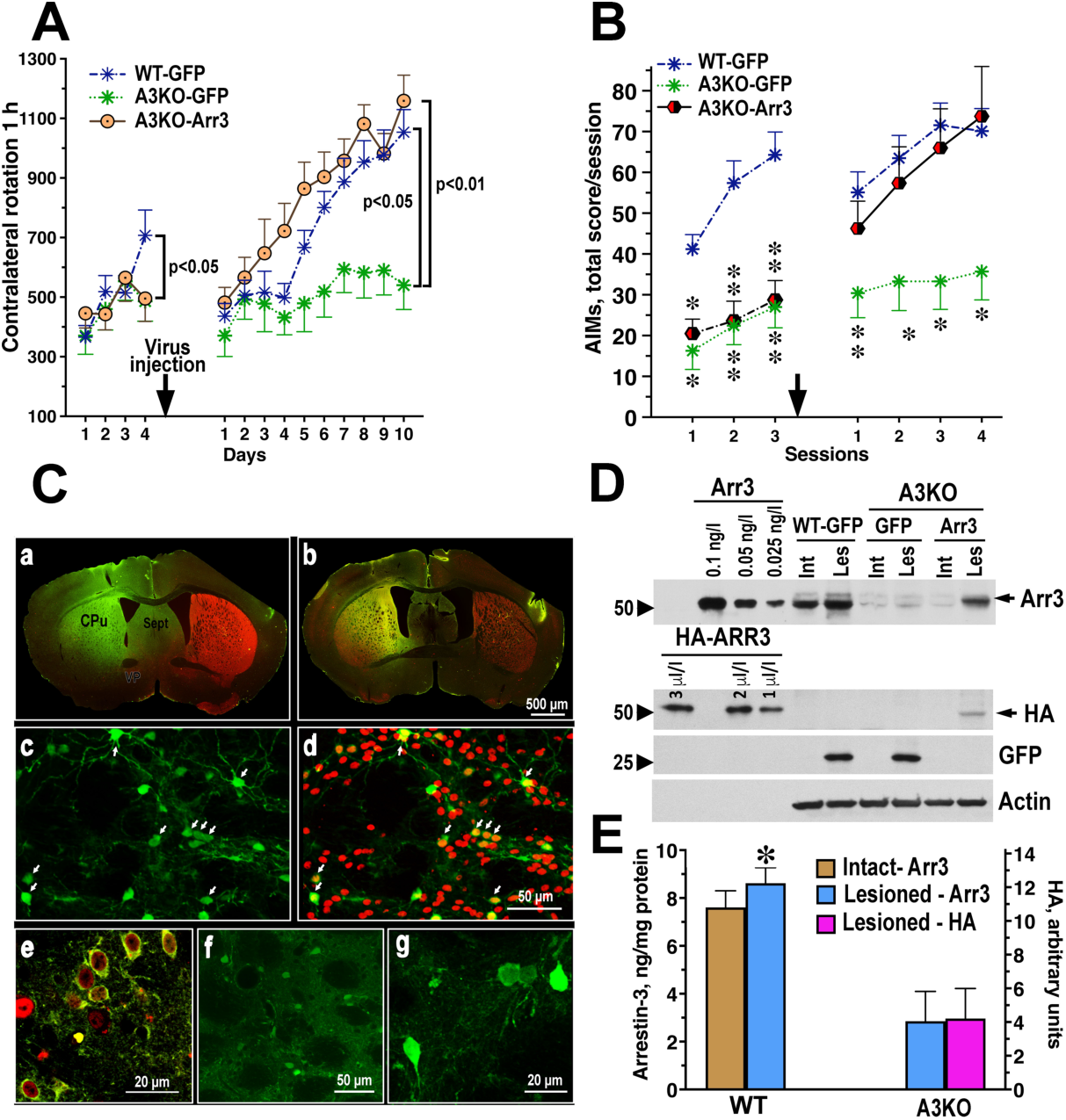
Lentivirus-mediated expression of Arr3 in the lesioned striatum rescues L-DOPA-induced rotations and AIMs in A3KO mice. (**A**) The indicated lentiviruses were injected into the dorsolateral striatum of A3KO mice. WT mice received GFP. WT Arr3 fully rescued rotations in A3KO mice. Significance by the Bonferroni post hoc test across all testing sessions is shown. (**B**) WT Arr3 fully rescues AIMs in A3KO mice. Arrow shows the time of the virus injection. * -p<0.05, ** -p<0.01 to WT for individual sessions by Dunn’s post hoc test following Kruskal-Wallis non-parametric ANOVA. (**C**) Immunohistochemical detection of protein expression in the lesioned striatum. (**a**) Low magnification photomicrograph of mouse brain. GFP (expressed co-cistronically with Arr3) in the lesioned striatum detected with mouse anti-GFP antibody (green) and TH detected with rabbit anti-TH antibody (red). (**b**) Low magnification photomicrograph of the mouse brain with Arr3 detected by co-cistronically expressed GFP (green) and a marker of MSNs FOXP1 (red). (**c,d**) WT Arr3 construct in MSNs labeled with GFP (green) alone (**c**) or co-localized with FOXP1 (red) (**d**). Arrows point to examples of co-labeled neurons in both images. (**e**) High magnification photomicrograph of the expression of WT Arr3 in MSNs co-labeled for GFP (green) and FOXP1 (red). (**f, g**) Low and high magnification photomicrographs of the expression of WT Arr3 labeled with anti-HA antibody (green). **(D)** <u>Upper panel</u>: Expression of WT Arr3 in the intact uninfected and lesioned infected striata of WT mice and A3KO mice infected with GFP or Arr3 lentivirus detected with anti-Arr3. The indicated amounts of purified Arr3 served as standards. Note the increased levels of endogenous Arr3 in the lesioned as compared to the intact striata. <u>Lower panel:</u> Expression of Arr3 in the lesioned hemisphere of A3KO mice infected with HA-Arr3 lentivirus detected with anti-HA antibody. Serial dilutions of lysates of HEK293 cells infected with HA-Arr3 lentivirus were used as standards to indicate the position of HA-Arr3 on the blot. Note the absence of HA-Arr3 in WT and A3KO mice infected with GFP and the presence of GFP in both. (**E**) Quantification of Western blot data. Known amounts of purified bovine Arr3 served as standards. Arr3 data are absolute numbers related to the endogenous Arr3 level in WT mice (≈35% of endogenous Arr3). The numbers for HA are arbitrary. Means±S.E.M. are shown. * -p<0.05 to the intact striatum by repeated measure ANOVA. N=8-13 mice per group.

We measured the rotation frequency in mice expressing GFP (control) or Arr3 on the lesioned side in the motor striatum. We first pre-tested the mice for rotations for 4 consecutive days (**Fig. 2A**). A3KO mice were then randomly assigned to 2 groups that received LVs encoding GFP or Arr3. A single group of WT mice expressing GFP served as an outgroup for comparison. During pre-testing, only the WT group showed sensitization (Genotype X Day p=0.0318 by repeated measure ANOVA across testing sessions). During the 10-day testing period following LV injection, A3KO mice expressing GFP displayed minimal sensitization, with the rotation frequency remaining essentially stable across testing sessions, whereas the expression of Arr3 fully rescued the rotational behavior (Genotype p=0.0022; Genotype X Day p=0.0019) (**Fig. 2A**). Next, we performed a similar rescue experiment using AIMs as a behavioral readout. During pretesting, A3KO mice demonstrated reduced AIMs compared to WT (p<0.001 by Mann-Whitney test by sessions). In A3KO mice expressing GFP, the level of AIMs remained low throughout the testing period. In contrast, the A3KO group expressing Arr3 demonstrated AIMs at a level similar to that of WT mice (group differences significant by Kruskal-Wallis test followed by Dann post hoc comparison, p<0.005) (**Fig. 2B**). Thus, Arr3 acts in the process of the behavioral sensitization.

Postmortem examination of the striatal samples demonstrated the expression of Arr3 in the MSNs of the lesioned striatum, as evidenced by co-staining with TH and FOXP1, a marker of MSNs (**Fig. 2C**), and the expression of Arr3 detected by Western blot with both anti-Arr3 and anti-HA antibodies (**Fig. 2D**). Quantification showed that LV-encoded Arr3 was expressed at ∼30-35% of the endogenous Arr3 level in WT mice (**Fig. 2E**).

### Arr3-mediated activation of the JNK pathway is required for its behavioral effect

Arrestins negatively regulate GPCR signaling via homologous desensitization (Gurevich and Gurevich, 2006). At the same time, arrestins facilitate signaling via other pathways (Gurevich and Gurevich, 2014; Gurevich and Gurevich, 2015, 2018b). The loss-of-function phenotype in A3KO mice suggests the loss of Arr3-mediated signaling. Arr3 is the only non-visual arrestin subtype that activates the JNK (McDonald et al., 2000; Seo et al., 2011; Song et al., 2009), so this pathway seemed a viable candidate. Initially it was reported that Arr3 selectively activated the JNK3 isoform (McDonald et al., 2000). We later found that Arr3 can activate at least some splice variants of JNK1 and JNK2 (Kook et al., 2014), although JNK3 is its preferred partner (Zhan et al., 2023; Zhan et al., 2014). To test the role of the JNK pathway in the behavioral function of Arr3, we used the Arr3-V343T mutant with impaired ability to activate JNK (Seo et al., 2011). As this was only demonstrated in non-neuronal cells, we compared the ability to facilitate JNK activation of WT Arr3 and the V343T mutant in human neuroblastoma SH-SY5Y cells and found that Arr3 increases JNK3 phosphorylation in these cells, whereas the V343T mutant has only minimal effect (**Fig. S1**). We compared the ability of Arr3-V343T and WT Arr3 to rescue L-DOPA-induced rotations in A3KO mice. We found that WT Arr3 afforded full rescue (p<0.001 to the GFP group), whereas Arr3-V343T failed to rescue rotations (**Fig. 3A**) despite comparable expression of both proteins (**Fig. 3B,C**).

**Figure 3.**
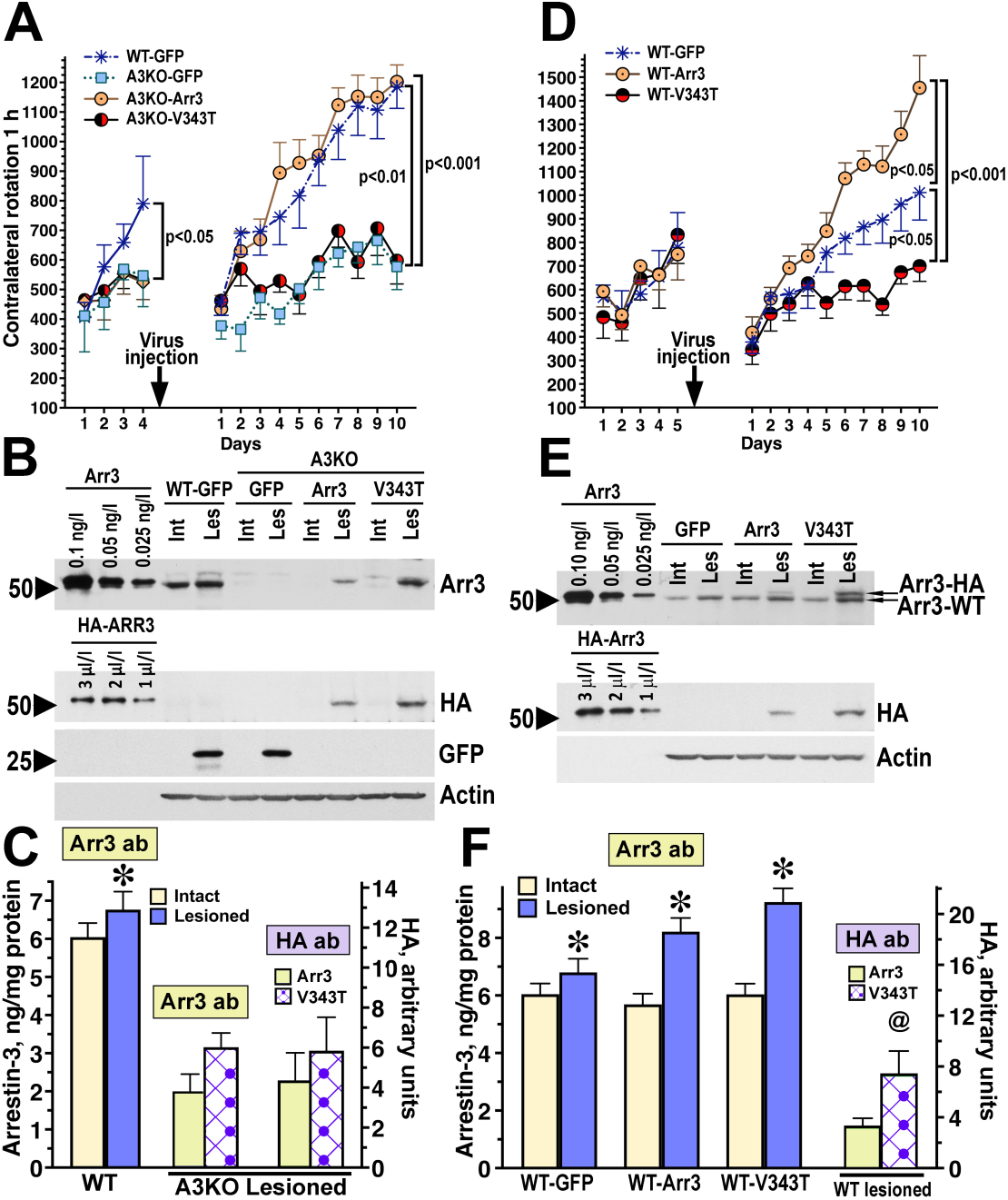
Modulation of L-DOPA-induced rotations by targeting Arr3-dependent JNK activation in WT and A3KO mice. **(A)** The indicated lentiviruses were injected into the dorsolateral striatum of A3KO mice. WT mice received GFP. Expression of WT Arr3 (Arr3) but not the V343T mutant rescued L-DOPA-induced rotations in A3KO mice. Significance shown by Bonferroni’s post hoc multiple comparison across testing days following two-way repeated measure ANOVA with GROUP and DAY as within group factors. For pre-injection testing, the significance value applies to comparisons between the WT-GFP group and each of the A3KO group. For the post-injection, the lower p value applies to the comparison between each of the GFP group to the A3KO-Arr3 and the large value – to the same comparisons with the WT-GFP group. **(B)** Expression of Arr3 WT and the V343T mutant in the intact uninfected and lesioned infected striata detected with anti-Arr3 (upper panel) and anti-HA (middle panel) antibody. Lysate of HEK293 cells infected with HA-Arr3 lentivirus was used to indicate the position of HA-Arr3 on the blot. GFP expression in control WT and A3KO groups and actin (loading control) are also shown. **(C)** Quantification of the Western blot data. Known amounts of purified bovine Arr3 (as shown in B, upper panel) were used as standards. * -p<0.05 to the value in the intact striatum by paired Student’s t-test. **(D)** Overexpression of WT Arr3 (Arr3), but not V343T, in WT mice increased the frequency of L-DOPA-induced rotations. V343T significantly reduced the rotation frequency in WT mice. The statistical analysis was the same as in A. **(E)** The anti-Arr3 antibody (upper panel) was used to detect both endogenous Arr3 (Arr3-WT) and overexpressed WT (Arr3) and mutant Arr3 (V343T). The anti-HA antibody detected the exogenous Arr3 constructs (middle panel). **(F)** Quantification of the Western blot data for the expression of the Arr3 and V343T constructs and endogenous Arr3 in the intact and lesioned hemisphere of WT mice infected with lentiviruses encoding Arr3 or V343T. The values for Arr3 in the lesioned injected hemisphere (blue bars) reflected the sum of endogenous and expressed Arr3 or endogenous Arr3 plus V343T. Means±S.E.M. are shown. N=10-13 mice per group. *, p<0.05 as compared to intact hemisphere by repeated measure ANOVA; @, p<0.05 to Arr3 by one-way ANOVA. See also **Fig. S1**.

Arr3-V343T can bind all components of the JNK pathway but fails to activate JNK3 (Seo et al., 2011). Therefore, we hypothesized that it could have a dominant-negative effect on JNK3 activity in the presence of endogenous Arr3. Indeed, testing the effect of WT Arr3 and V343T in WT mice we found that WT Arr3 further enhanced the rotation frequency (p<0.05) (**Fig. 3D**), as compared to GFP control. In contrast, the V343T mutant significantly reduced the rotation frequency across testing days (p<0.05 to WT-GFP). The behavior of WT mice expressing V343T resembled that of A3KO mice. We detected significantly higher expression of V343T than of the WT Arr3, as measured with anti-HA antibody (**Fig. 3E,F**), although the expression of both WT and V343T Arr3 was relatively low in comparison to the level of endogenous Arr3 (**Fig. 3F**).

### Arr3-mediated activation of JNK in the direct pathway MSNs is sufficient to produce the pro-sensitization effect

The data with the V343T mutant suggested that Arr3-dependent JNK activation plays a key role in its pro-sensitization effect. However, Arr3 is a multifunctional protein, and other functions might have been altered by the V343T mutation. To specifically test the role of Arr3-dependent JNK activation, we used the monofunctional Arr3-derived peptide T1A that acts as a mini-scaffold facilitating JNK activation in cells (Perry-Hauser et al., 2022; Zhan et al., 2016). As this has been shown only in non-neuronal cells (Perry-Hauser et al., 2022; Zhan et al., 2016), first we ascertained that Venus-tagged T1A facilitated JNK3 activation in human neuroblastoma SH-SY5Y cells, whereas the homologous arrestin-2-derived peptide B1A did not (**Fig. S2A,B**). The expression of Venus-T1A in the lesioned striata of A3KO mice rescued L-DOPA-induced rotations as effectively as full-length Arr3 (p<0.001 to both A3KO-GFP and B1A groups) (**Fig. 4A**). In contrast, Venus-B1A, a peptide homologous to T1A derived from arrestin-2, which does not activate JNK (Miller et al., 2001; Seo et al., 2011)(**Fig. S2A,B**), was ineffective (**Fig. 4A**), despite comparable expression of Venus-T1A and Venus-B1A (**Fig. 4C,D**).

**Figure 4.**
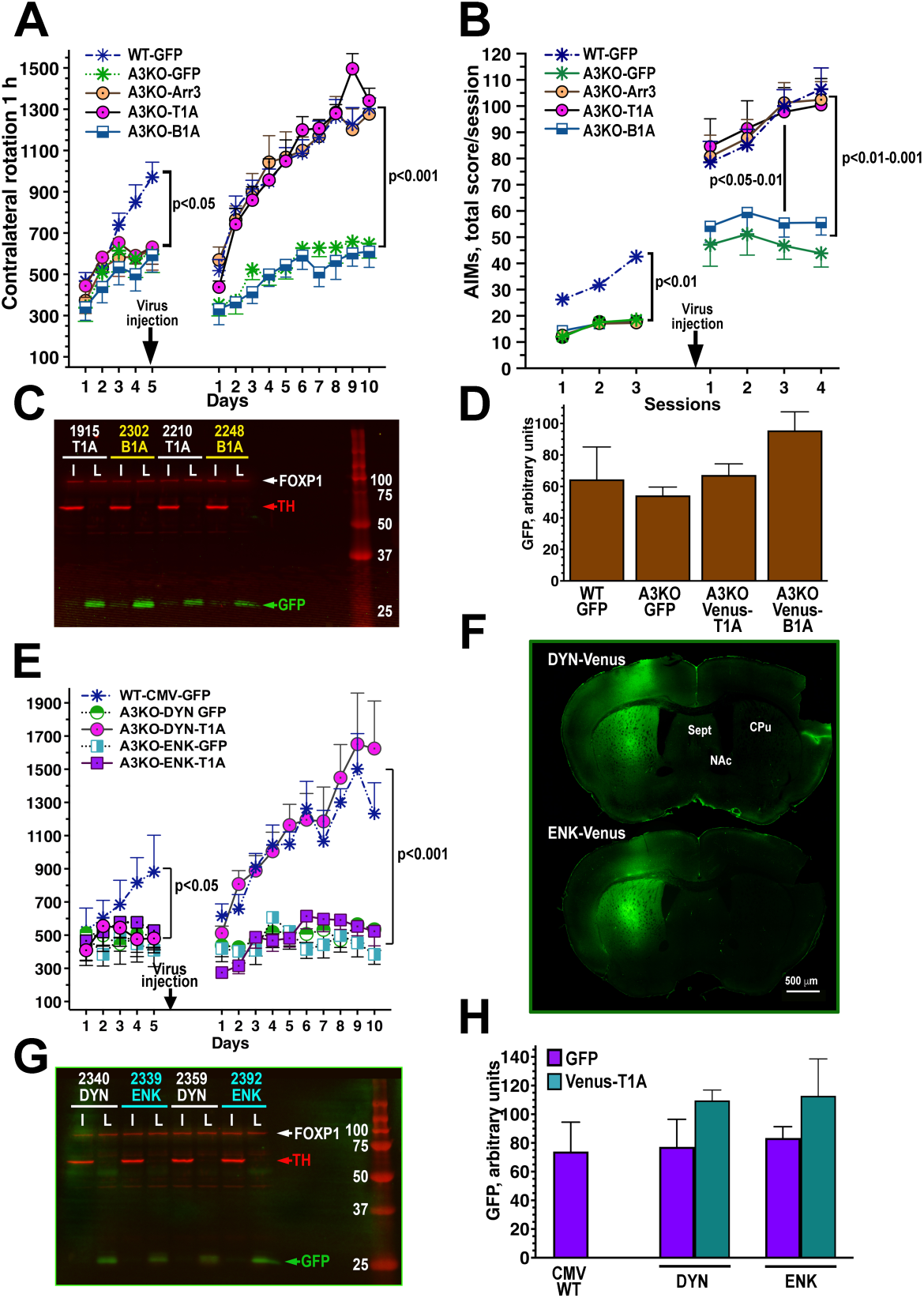
Rescue of L-DOPA-induced rotations and AIMs in A3KO mice by the JNK3-activating Arr3-derived peptide T1A. **(A)** Both WT Arr3 and the Arr3-derived peptide T1A fully rescued L-DOPA-induced rotations in A3KO mice. Arrestin-2-derived homologous B1A peptide failed to rescue rotations. Significance shown after the virus injection applies to pairwise comparisons between WT-GFP and A3KO-T1A with A3KO-GFP and A3KO-B1A by Bonferroni’s post hoc comparison following two-way repeated measure ANOVA. Before virus injection, WT was the only group significantly different from the rest. **(B)** WT Arr3 and T1A fully rescued AIMs in A3KO mice whereas B1A did not. The data were analyzed by Dunn’s post hoc test following Kruskal-Wallis non-parametric ANOVA for each session. Before the virus injection, the WT group was the only one significantly different from the rest. Following the injection, the A3KO groups expressing Arr3 and T1A showed full rescue of the AIMs behaviors to the level of WT, whereas A3KO groups expressing GFP and B1A remained significantly below. Significance levels shown apply to pairwise comparisons between each of the group WT, A3KO/Arr3, and A3KO/T1A versus each of A3KO/GFP and A3KO/B1A (smaller p value applies to GFP). The differences in Sessions I and II are the same as in Session III (not shown for lack of space). **(C)** Western blot detection of the expression of Venus-T1A and Venus-B1A with mouse anti-GFP antibody followed by anti-mouse IRDye 800CW antibody. Tyrosine hydroxylase and a marker of MSNs FOXP1 were detected with the respective rabbit antibodies followed by anti-rabbit IRDye 680RD secondary antibodies. The blots were visualized with the Odyssey CLX imaging system. **(D)** Quantification of the Western blot data for the expression of Venus-T1A and Venus-B1A in the lesioned hemisphere detected with the anti-GFP antibody. The expression of GFP in control GFP-injected groups of WT and A3KO mice is shown for comparison. There were no significant differences in the expression of the Venus-fused peptides among the experimental groups as revealed by one-way ANOVA (p=0.22). (**E)** The Arr3-derived peptide T1A, selectively expressed in striatonigral MSNs under the control of the DYN promoter, fully rescued rotations in A3KO mice. In contrast, selective expression of T1A in striatopallidal MSNs under the control of the ENK promoter was ineffective. Significance is by the Bonferroni post hoc test across all testing sessions. Significance shown after the virus injection applies to pairwise comparisons between WT-GFP and A3KO-DYN-T1A with A3KO-DYN-GFP, A3KO-ENK-GFP and A3KO-ENK-T1A. Before the virus injection, WT was the only group significantly different from the rest. **(F)** Images of the mouse brain stained for GFP demonstrating the level of expression driven by the DYN promoter as compared to the ENK promoter. Spt, septum; NAC, nucleus accumbens; CPu, caudatoputamen. **(G)** The expression of Venus-T1A driven by the DYN and ENK promoters detected by Western blot with mouse anti-GFP using the Odyssey imaging system. **(H)** Quantification of the Western blot data for the expression of Venus-T1A in the lesioned hemisphere driven by the DYN or ENK promoter detected with the anti-GFP antibody. The expression of GFP in control GFP-injected groups of WT mice is shown for comparison. There were no significant differences in the expression level of Venus-T1A among the experimental groups (p=0.43 one-way ANOVA). N=8-9 mice per group. N=9-11 mice per group. See **also Figs. S2**, **S3** and **S4**. See also **Videos S1** and **S2**.

We also performed a similar rescue experiment using AIMs as a behavioral readout. During pretesting, A3KO mice demonstrated reduced AIMs compared to WT (p<0.001 by Mann-Whitney test by sessions). In A3KO mice expressing GFP, the level of AIMs remained low throughout the testing period. In contrast, the A3KO group expressing WT Arr3 demonstrated AIMs at a level similar to that of WT mice (group differences significant by Kruskal-Wallis test, p<0.005) (**Fig. 2B**). Venus-T1A rescued AIMs as effectively as full-length Arr3 (**Fig. 4B**). The analysis of individual AIMs scores (**Fig. S3A-C, also see Videos S1 and S2**) supported the conclusion that the lack of Arr3 suppressed behavioral sensitization to L-DOPA, particularly evidenced by the reduction of the frequency of locomotor and axial AIMs (**Fig. S3B,C**), whereas the level of these behaviors remained high in the groups expressing Arr3 or T1A. Thus, Arr3 plays a direct role in behavioral sensitization via JNK activation.

The direct and indirect pathway MSNs differ in the expression of the dopamine receptors and neuropeptides, as well as in their role in movement control (Gerfen, 2022; Keeler et al., 2014). The two non-visual arrestins are equally expressed in both types of MSNs (Bychkov et al., 2012). To determine the site of Arr3 action in these behavioral paradigms, we constructed LVs with Venus-T1A under the control of dynorphin (DYN) or encephalin (ENK) promoters to target the expression specifically to the direct or indirect pathway neurons, respectively. To increase expression, Venus-T1A was placed under the additional control of the internal ribosome entry site (IRES), termed superIRES for its ability to strongly enhance translation (Bochkov and Palmenberg, 2006) (**Fig. S4A**). These vectors yielded expression comparable to that driven by the strong CMV promoter (**Fig. S4B**). Although these promoters have been used previously (Ferguson et al., 2011), we confirmed their neuronal selectivity in the mouse brain (**Fig. S4C**).

The expression of Venus-T1A under the control of the ENK promoter was ineffective in restoring rotational behavior in A3KO mice. Mice expressing ENK-Venus-T1A did not differ from mice expressing GFP under the control of the ENK promoter. In contrast, when Venus-T1A was expressed under the DYN promoter, a complete rescue was observed (p<0.001 to the DYN-GFP group) (**Fig. 4E**). Both promoters yielded comparable expression in the brain, as evidenced by immunohistochemistry (**Fig. 4F**) and Western blot of samples collected postmortem (**Fig. 4G,H**). Thus, Arr3-dependent JNK activation specifically in the direct pathway neurons critically contributes to the dopaminergic behavioral sensitization.

### Signaling mechanisms and JNK3 activity in the lesioned striatum

To gain insight into the mechanisms of Arr3 action in behavioral sensitization, we examined the activity of signaling pathways known to be altered by persistent treatment with dopaminergic drugs (Ahmed et al., 2010; Ahmed et al., 2015a; Bastide et al., 2015; Beaulieu et al., 2005; Bychkov et al., 2007). The lesion-induced super-sensitivity of the ERK pathway has been implicated in the so-called priming process associated with behavioral sensitization (Ahmed et al., 2015a; Bastide et al., 2015; Bychkov et al., 2007). Both arrestin-2 and Arr3 have been shown to scaffold the ERK cascade facilitating ERK1/2 activation (Luttrell et al., 2001). We detected the expected super-responsiveness of ERK1/2 to L-DOPA challenge in the lesioned striatum, but no difference between WT and A3KO mice (**Fig S5A,B**). Similarly, the Akt pathway in the DA-depleted striatum has been shown to become constitutively supersensitive following chronic L-DOPA treatment (Ahmed et al., 2015a; Bychkov et al., 2007). The degree of Akt supersensitivity was also similar in WT and A3KO mice (**Fig S5E-G).** There is no direct evidence that arrestins activate another MAP kinase pathway, p38, although the p38 kinase, like JNK, is activated by a variety of cellular stresses and shares many upstream kinases with the JNK pathway (Ono and Han, 2000). We have previously shown that the p38 pathway is activated by L-DOPA challenge in both intact and lesioned striata (Ahmed et al., 2015a; Bychkov et al., 2007). The degree of p38 activation by L-DOPA was similar in WT and A3KO mice (**Fig. S5C,D**).

We also examined the activity of the JNK-c-Jun pathway in the lesioned striatum of WT and A3KO mice. Ten distinct JNK isoforms are expressed in the brain: four splice variants of JNK1, four of JNK2, and two of JNK3 (**Fig. S6A**). Phospho-JNK (ppJNK) is seen as two major bands at 54 and 46 kDa (**Fig. S6B**), which contain nine out of the ten known JNK isoforms (Coffey, 2014). In the mouse striatum, in contrast to the rat, the longer JNK3 splice variant JNK3α2 (57 kDa) is undetectable when probed with a ppJNK-specific antibody (**Fig. S6B**). The shorter JNK3α1 is the major splice variant in both rat and mouse striata. This isoform runs at 54 kDa together with two splice variants of JNK2 (JNK2α2 and JNK2β2) and two splice variants of JNK1 (**Fig. S6B**). The lower band contains two shorter JNK2 and two JNK1 splice variants.

We specifically tested JNK3 activity in the striata of WT and A3KO mice. The anti-JNK3 antibody precipitated both JNK3 isoforms and the samples were then immunoblotted for phospho-JNK (**Fig. 5A,D**). We compared the fraction of active (phosphorylated normalized to total immunoprecipitated) JNK3 in the intact and lesioned striata of WT and A3KO mice chronically treated with saline or L-DOPA and challenged with L-DOPA 45 min before sacrifice. We found no differences between the genotypes regardless of the treatment in the intact hemisphere (**Fig. 5A-C**). In contrast, in the lesioned hemisphere WT mice had a significantly higher activity of both isoforms of JNK3, particularly of JNK3α2, than A3KO mice (**Fig. 5A-C**). To test the specific effect of Arr3 on the JNK3 activity in the lesioned striatum, we compared the activity of the JNK3 isoforms in WT mice and A3KO mice injected with LVs encoding GFP (control) or WT Arr3. We found that, in parallel with the behavioral sensitization, exogenous Arr3 rescued JNK3 activation in A3KO mice (**Fig. 5D-F**).

**Figure 5.**
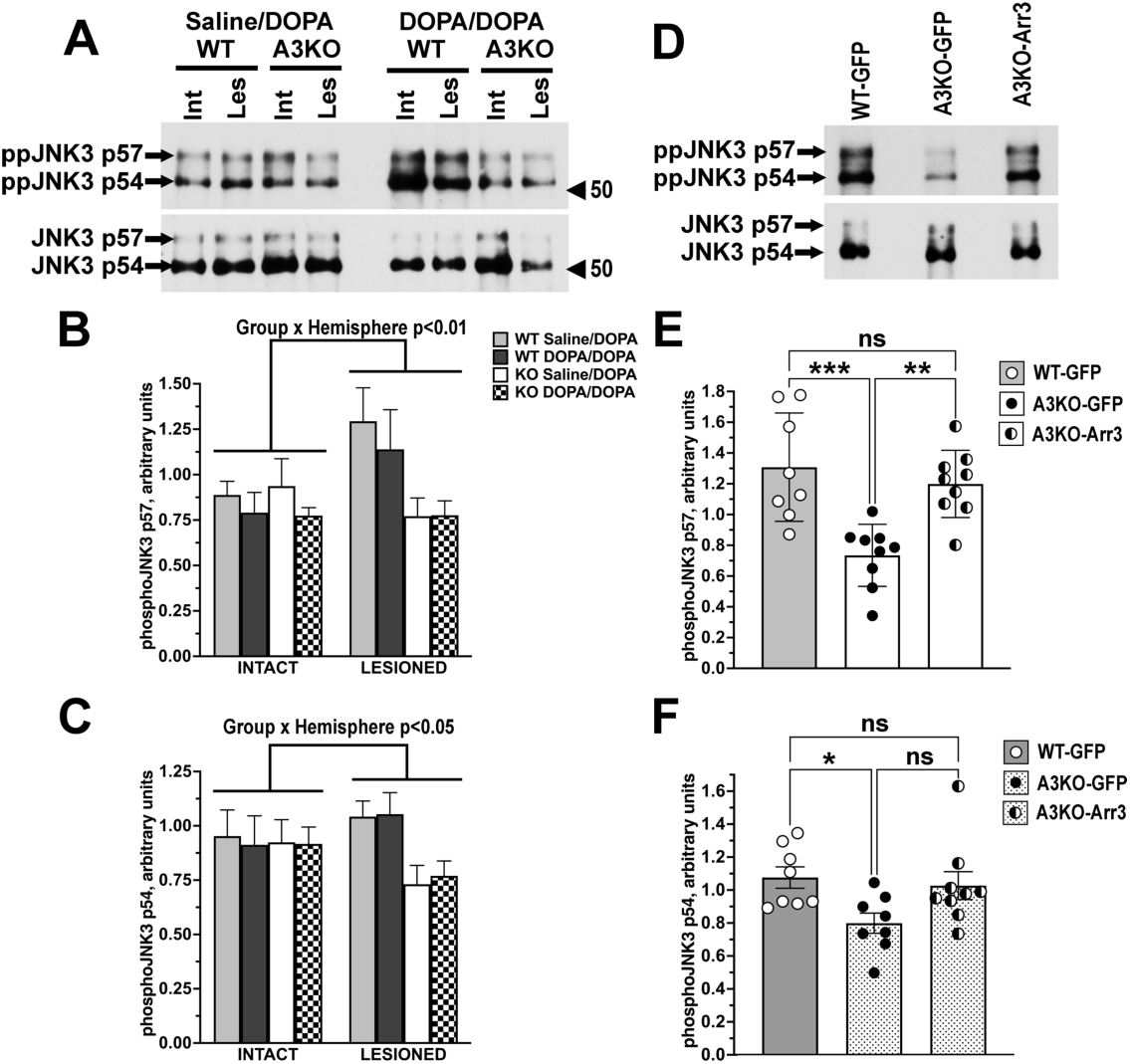
Changes in the activity of JNK3 in the striatum associated with the loss of Arr3. **(A)** Representative Western blots showing phosphorylated JNK3 isoforms (upper panel) and the total JNK3 (lower panel) in the intact and lesioned striata of WT and A3KO mice. JNK3 was immunoprecipitated from the intact and lesioned striata of WT and A3KO mice chronically treated with either saline or L-DOPA for 10 days and challenged with L-DOPA as described in Methods. Arrows and numbers point to JNK isoforms. **(B)** Quantification of the Western blot data for phospho-JNK p57 isoform. Note that the value for phospho-JNK3 p57 was normalized for immunoprecipitated JNK3 p57 in each sample. The two-way repeated measure ANOVA with GROUP as between group and Hemisphere as within group factors yielded significant effect of Hemisphere and significant Group x Hemisphere interaction (p<0.01), which is due to enhanced activation of JNK3 p57 in the lesioned hemisphere of WT mice. If the analysis is restricted to the lesioned hemisphere, the level of JNK3 p57 phosphorylation in A3KO mice is significantly lower than in WT mice (p<0.05). N = 5-7. **(C)** Quantification of the Western blot data for phospho-JNK p54 isoform. The same statistical analysis as for p57 yielded significant Group x Hemisphere interaction (p<0.05), which is due to enhanced activation of JNK3 p54 in the lesioned hemisphere of WT mice. **(D)** Representative Western blots showing phosphorylated JNK3 isoforms (upper panel) and the total JNK3 (lower panel) in the lesioned striata of WT mice injected with the GFP lentivirus and A3KO mice injected with either GFP or Arr3 lentivirus. **(E)** Quantification of the Western blot data for phospho-JNK p57 isoform. One-way ANOVA analysis yielded highly significant effect of Group (p=0.0003). The results of subsequent post-hoc comparisons by Tukey’s multiple comparison test are shown on the graph. ** p<0.01, *** -p<0.001. **(F)** Quantification of the Western blot data for phospho-JNK p54 isoform. The effect of Group was significant (P=0.0324). The results of subsequent post-hoc comparisons by Tukey’s multiple comparison test are shown on the graph. * -p<0.05.

The transcription factor c-Jun is the best-known substrate of JNKs (Bogoyevitch and Bostjan Kobe, 2006), which gave this family of kinases its name (c-<u>J</u>un <u>N</u>-terminal <u>K</u>inase). JNK-dependent phosphorylation and activation of c-Jun has been extensively studied in the context of the JNK role in apoptosis (Coffey, 2014; Zhan et al., 2014). The role of JNK-c-Jun signaling in neural processes unrelated to cell death is beginning to be appreciated (Coffey, 2014; Hollos et al., 2018). We examined the level of c-Jun phosphorylation in the brains of WT and A3KO mice, both drug-naïve animals challenged with L-DOPA and mice chronically treated with L-DOPA. Chronic L-DOPA administration significantly increased the JNK-dependent c-Jun phosphorylation in the dopamine-depleted striatum in L-DOPA-treated WT, but not in A3KO mice (**Fig. 6A-C**). There was a statistically significant reduction in the concentration of phospho-c-Jun (**Fig. 6C,D**) (p<0.01) in the lesioned striatum of chronically L-DOPA-treated A3KO animals as compared to WT. The LV-mediated expression of Arr3 rescued c-Jun phosphorylation (**Fig.6D-F**) Thus, the activity of the JNK pathway in the striatum of A3KO mice is reduced, resulting in diminished JNK-dependent phosphorylation of c-Jun.

**Figure 6.**
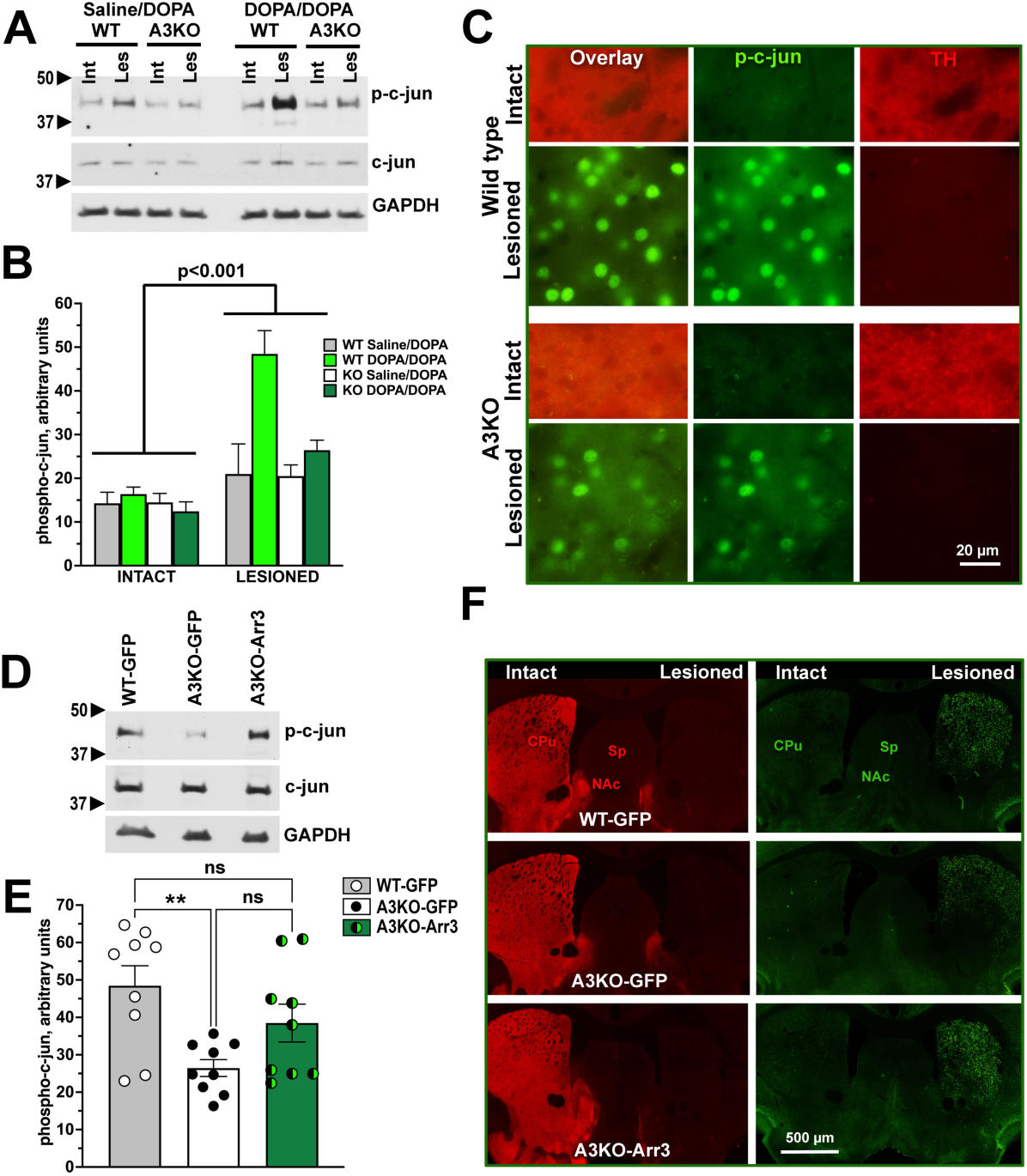
Changes in the JNK-dependent c-Jun activation associated with the loss of Arr3. **(A)** Representative Western blots showing the levels of c-Jun phosphorylated at the JNK site Ser73 (upper panel) and total c-Jun (middle panel) in the intact and lesioned striata of drug-naïve or chronically L-DOPA-treated WT and A3KO mice. Numbers on the left are molecular weight standards. **(B)** Quantification of the Western blot data for phospho-c-Jun in the intact and lesioned striata in WT and A3KO mice. Highly significant Hemisphere x Group interaction (p<0.0001) is due to elevated c-Jun phosphorylation in the lesioned striatum of L-DOPA-treated WT mice, which is not evident in other groups. N=6-9. (**C**) High power images of the mouse brain double-stained for phospho-c-Jun (green) and TH (red) demonstrating elevated phospho-c-Jun in the lesioned hemisphere and reduced c-Jun phosphorylation in the lesioned striatum of A3KO mice. **(D**) Representative Western blots showing the levels of c-Jun phosphorylated at the JNK site in the lesioned striatum of L-DOPA-treated WT mice expressing GFP and A3KO mice expressing either GFP or Arr3 (A3KO-Arr3). (**E)** Quantification of the Western blot data for phospho-c-jun. The data analyzed by one-way ANOVA (p=0.007) followed by post-hoc comparisons by Tukey’s multiple comparison test show significantly reduced level of phospho-c-Jun in GFP-expressing A3KO mice as compared to both WT-GFP. The values in the A3KO-Arr3 did not differ significantly from either WT-GFP or A3KO-GFP group. ** -p<0.01 **(F)** Low power photomicrographs of mouse brain sections co-stained for TH (red) and phospho-c-Jun (green) showing reduced level of phospho-c-Jun in A3KO-GFP mice rescued by expression of Arr3.

Our data suggest that an increased level of c-Jun phosphorylation dependent on Arr3-assisted JNK activation is necessary for the development of behavioral sensitization. A3KO mice display diminished sensitization because of the absence of this mechanism.

### Arr3 regulates behavior via scaffolding of the JNK activation cascade

It has been reported previously that Arr3 suppresses behavioral sensitization when overexpressed in the lesioned striatum of mice and non-human primates (Urs et al., 2015). Our results in A3KO mice appear to contradict this. The previous study utilized adeno-associated viruses (AAVs) as vectors. AAVs are known to induce high expression levels. The JNKs are the output kinases activated by a three-tiered cascade of kinases that sequentially phosphorylate and activate the downstream kinase (Weston and Davis, 2007). We previously demonstrated that Arr3 facilitates activation of JNKs via simple scaffolding. That was evidenced by the bell-shaped curve of the dependence of the JNK3 phosphorylation on Arr3 in non-neuronal cultured cells (Kook et al., 2014; Zhan et al., 2023; Zhan et al., 2011; Zhan et al., 2013). As demonstrated *in silico*, a bell-shaped curve is characteristic for the simple scaffolding mechanism: while low scaffold concentrations promote activation, high concentrations beyond certain point invariably decrease the output (Levchenko et al., 2000). Here we confirmed the scaffolding nature of the Arr3-mediated JNK3 activation in neuronal SH-SY5Y cells (**Fig. S7**). Thus, the effect of Arr3, acting as a scaffold *in vivo*, on the JNK activity and behavior likely depends on its concentration. Therefore, we tested the effect of Arr3 on the dopaminergic behavioral plasticity as a function of its expression level. To this end, we compared the effects of the Arr3 gene transfer mediated by AAV and LV on the rotational and AIMs sensitization in WT mice. In this experiment, we confirmed that the LV-mediated expression of Arr3 in the lesioned striatum of WT mice facilitated locomotor sensitization to L-DOPA (**Fig. 3D**). In contrast, the AAV-mediated expression of Arr3 significantly suppressed sensitization of the rotational behavior (**Fig. 7A**). The LV and AAV-mediated Arr3 expression in the lesioned striata also produced opposite effects on the frequency of AIMs (**Fig. 7B**). Comparison of the level of Arr3 expression confirmed that AAV-driven expression was much higher that the LV-driven expression (**Fig. 7C-E**). The LV-driven expression was at 30-35% of the endogenous level (**Fig. 8C,D**), which is consistent with the data from previous experiments (**Fig. 2E, 3C,F**). In contrast, the AAV-mediated expression exceeded the endogenous Arr3 level in WT mice approximately 10-fold (**Fig. 7D**).

**Figure 7.**
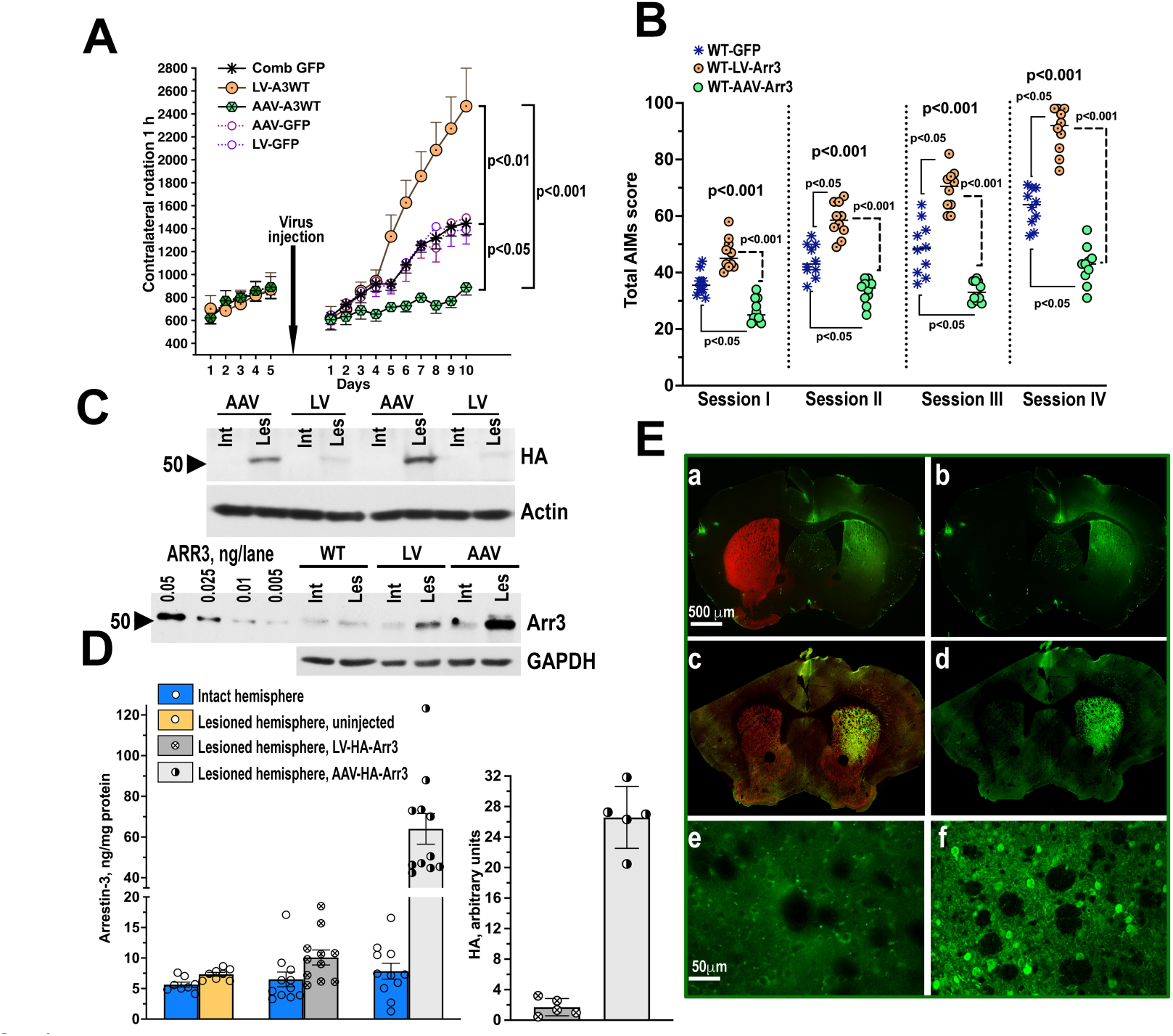
Modulation of L-DOPA-induced rotations by the LV- or AAV-mediated Arr3 expression in the striatum of WT mice. **(A)** The indicated viruses were injected into the dorsolateral striatum of WT mice. The control group receive GFP via AAV. The expression of WT Arr3 driven by LV enhanced L-DOPA-induced rotations, whereas the AAV-mediated expression suppressed sensitization of rotational behavior (two-way repeated measure ANOVA, Genotype effect F(2,270)=17.9 p<0.001; Genotype X Day effect p<0.001); the significance shown is by Bonferroni across days. **(B)** Similarly, the expression of WT Arr3 driven by LV enhanced L-DOPA-induced AIMs, whereas the AAV-mediated expression suppressed AIMs, resulting in no sensitization of these behaviors. The graph shows results for each post-injection testing session analyzed separately by Kruskal-Wallis non-parametric ANOVA (significance levels shown above). The brackets indicate significant difference to WT-GFP group by post-hoc Dunn’s multiple comparisons. **(C)** <u>Upper panel:</u> Expression of HA-tagged Arr3 WT mediated by LV or AAV in the intact uninfected and lesioned infected striata detected with anti-HA antibody. Actin shown as loading control. <u>Lower panel:</u> Expression of HA-tagged Arr3 mediated by LV or AAV in the intact uninfected and lesioned infected striata detected with anti-Arr3 antibody. Known amounts of purified Arr3 (four left lanes) served as standards for quantification. The load for the AAV-injected sample is 1/3 of the rest to allow for visual presentation on one blot. Note that endogenous Arr3 and exogenous HA-Arr3 are not resolved on the gel. GAPDH is shown as loading control. **(D)** Quantification of the Western blot data. Scatterplots with means±S.E.D. are shown. The values obtained with anti-Arr3 antibody were converted into absolute numbers (ng Arr3/mg total protein) using known amounts of purified bovine Arr3 as standards to allow for comparison with the level of endogenous Arr3 in the intact and lesioned hemisphere of WT mice. Values obtained with anti-HA antibody are expressed in arbitrary units. **(E)** Images of the mouse brain stained for GFP demonstrating the level of expression driven by LV and AAV. **a,b** - a representative striatal section from a mouse following LV-Arr3 injection co-stained for GFP and TH; **a** - an overlay, **b** - green channel for GFP; **c,d** - a representative striatal section from a mouse following AAV-Arr3 injection co-stained for GFP and FOXP1; **c** - an overlay, **d** - green channel for GFP; **e,f** - high power photomicrographs of the sections from the striatum infected with LV-HA-Arr3 **(e)** or AAV-HA-Arr3 **(f)** stained for HA. Abbreviations as in Fig. 5C.

## DISCUSSION

Our main finding is that the mice lacking one of the ubiquitous arrestin isoforms, Arr3, but not arrestin-2, following unilateral DA depletion display a reduced propensity for behavioral sensitization by chronic treatment with the dopamine precursor L-DOPA. The behavioral sensitization was measured as increased responsiveness to the drug in two independent tests, contralateral rotations and AIMs. Contralateral rotations are a behavioral response to high doses of L-DOPA and many DA agonists in mice with unilaterally ablated dopaminergic neurons in the substantia nigra that provide dopamine to the striatum. The frequency of these rotations progressively increases upon chronic treatment (Ahmed et al., 2010; Ahmed et al., 2015a; Ahmed et al., 2015b; Ahmed et al., 2007; Bychkov et al., 2007). Lower L-DOPA doses bring about a collection of orofacial, locomotor, and trunk movements collectively known as <u>A</u>bnormal Involuntary <u>M</u>ovements (AIMs) (Andreoli et al., 2021; Cenci and Lundblad, 2007), the frequency of which also increases with each L-DOPA administration. The phenomenon of progressively increasing behavioral responsiveness to repeated drug administration is referred to as behavioral sensitization. We demonstrated that Arr3 is required for sensitization, since A3KO mice do not manifest it, whereas exogenous Arr3 supplied to the striatum via viral gene transfer fully rescued the sensitization of both behaviors in A3KO mice.

We found that Arr3 contributes to the sensitization process via Arr3-assisted activation of the JNK pathway, specifically the JNK3 activation in the direct pathway striatal MSNs. This conclusion is based on the failure of an Arr3 mutant Arr3-V343T defective in the JNK activation to rescue sensitization, as well as on its ability to inhibit sensitization in WT animals, apparently acting in a dominant-negative manner. Sensitization of both behaviors was also rescued in A3KO mice by an Arr3-derived short peptide T1A capable of activating JNK but lacking other functions of the full-length Arr3 protein (Perry-Hauser et al., 2022; Zhan et al., 2016). Additional support for the role of the JNK pathway is provided by the expected supersensitivity in A3KO mice of other signaling pathways previously found to be abnormal following DA depletion and/or L-DOPA treatment in WT animals [(Ahmed et al., 2010; Ahmed et al., 2015a; Ahmed et al., 2007; Bochkov and Palmenberg, 2006); see also (Bastide et al., 2015) and references therein], along with defective JNK3 activation in A3KO mice, which is rescued by virally-delivered Arr3 in parallel with the behavioral rescue. By demonstrating that an Arr3-derived JNK-activating peptide T1A incapable of interacting with GPCRs fully rescues L-DOPA-induced behavioral sensitization, we ruled out the role of the Arr3 action at GPCRs in the sensitization mechanism. Our data implicate Arr3-dependent JNK3 activation in the L-DOPA-induced behavioral plasticity.

Arrestins are known to regulate a multitude of signaling pathways (Gurevich and Gurevich, 2023; Peterson and Luttrell, 2017; Wess et al., 2023), with arrestin-2 and Arr3 interacting with >100 proteins each (Xiao et al., 2007). This made identifying the specific pathway(s) involved in a physiological and/or behavioral response a challenge. This is one of very few studies focused on arrestin-mediated signaling where the exact arrestin-regulated pathway responsible for the physiological response in living animals has been identified (Beaulieu et al., 2005; Urs et al., 2011). It is also the first study implicating the JNK pathway in behavioral sensitization to L-DOPA. The activity of the JNK pathway in general and JNK3 in particular is most often regarded in the context of cell death in neurodegenerative disorders (Busquets et al., 2019; Hepp Rehfeldt et al., 2020; Musi et al., 2020). However, JNK3 has functions in the brain unrelated to neuronal death (Coffey, 2014; Musi et al., 2020; Nakano et al., 2020; Priori et al., 2023).

Our data suggest that Arr3 acts in the direct pathway MSNs expressing the D1 DA receptor. Numerous studies have demonstrated striking supersensitivity of D1 receptors brought about by DA depletion, which may or may not be ameliorated by subsequent L-DOPA treatment [(Ahmed et al., 2010; Aubert et al., 2005; Gerfen, 2000; Gerfen et al., 2002); see also (Bastide et al., 2015) and references therein]. The studies of the behavioral manifestations caused by dopaminergic drugs in animals with DA depletion have focused on GPCRs expressed by MSNs, primarily DA receptors. Since arrestins were initially identified as GPCR-binding proteins, the arrestin-mediated signaling was also originally envisioned as the function of an arrestin bound to a GPCR (Lefkowitz and Shenoy, 2005; Lefkowitz and Whalen, 2004). This is perfectly correct for the arrestin-dependent regulation of ERK1/2, as only arrestins bound to activated phosphorylated GPCRs have high affinity for ERK (Luttrell et al., 2001; Song et al., 2009) and facilitate ERK1/2 activation (Breitman et al., 2012; Luttrell et al., 2001). In contrast, the JNK pathway is regulated by a non-receptor bound Arr3 (Breitman et al., 2012; Miller et al., 2001; Song et al., 2009; Zhan et al., 2016; Zheng et al., 2023). This leaves open the question about a link between the stimulation of DA receptors and Arr3-dependent JNK3 activation. The dopaminergic activity is indispensable, since without L-DOPA no sensitization is observed. It is tempting to speculate that one or more of the upstream MAP kinase kinase kinases (MAP3K) serves as a link (Cuevas et al., 2007; Gurevich and Gurevich, 2018a). MAP3Ks are referred to as “signaling hubs” that integrate a multitude of stimuli and provide the selectivity and spatio-temporal control over the MAP kinase signaling (Cuevas et al., 2007). There is so far no evidence that MAP3Ks are directly activated by D1 receptor stimulation, but indirect activation by cellular stress caused by the high level of DA upon L-DOPA administration could be envisioned. Thus, the activation of one or more MAP3Ks upstream of the JNK pathway might serve as a link between the DA D1 receptor stimulation and Arr3-mediated JNK3 activation. This issue awaits further investigation.

Arrestins serve as scaffolds of the three-tiered MAP kinase activation cascades (Luttrell et al., 2001; McDonald et al., 2000; Peterson and Luttrell, 2017). In mechanistic terms, scaffolding means that arrestins, lacking their own enzymatic activity, facilitate signaling via an assembly of multi-protein complexes, bringing the kinases into close proximity and thus promoting their sequential activation. Mathematical modeling has revealed a biphasic dependence of the signaling effect on the scaffold concentration, with a lower concentration enhancing and a higher concentration inhibiting the signaling (Levchenko et al., 2000; Levchenko et al., 2004). Mechanistically, this is intuitive: when there are fewer molecules of the scaffold than of the scaffolded kinases, its presence increases the probability of the assembly of complete three-kinase signaling modules, whereas when there is more scaffold than kinases, mostly incomplete unproductive complexes are formed. In fact, one of the scaffolds of the JNK pathway, JIP1, was first discovered as a suppressor of JNK signaling (Dickens et al., 1997). We have previously shown that Arr3 activates the JNK pathway by acting as a scaffold (Kook et al., 2014; Zhan et al., 2011; Zhan et al., 2013). Therefore, our finding that low and high Arr3 expression levels have opposite effects on the behavioral sensitization in living mice, with a low Arr3 level promoting and a high level inhibiting sensitization in both behavioral paradigms, is not surprising. The most parsimonious explanation is that Arr3 acts as a scaffold for the JNK pathway in striatal neurons *in vivo* as it was shown to do in cultured cells *in vitro*. Our data explain earlier published results demonstrating that AAV-mediated overexpression of Arr3 in the lesioned striatum of WT hemiparkinsonian rodents suppressed AIMs (Urs et al., 2015). Although no quantification of the Arr3 expression levels was provided in that study (Urs et al., 2015), it is well known that AAV-mediated gene transfer usually yields high expression. We found the LV-mediated expression to be in the range of 25-30% of the endogenous Arr3 level, whereas the AAV-mediated Arr3 expression exceeded the WT level ∼10-fold. Thus, our data provide the first experimental evidence of the Arr3 scaffolding action for the JNK activation cascade in living animals.

The unilaterally 6-OHDA lesioned rodents have been extensively used as an animal model of LID in PD (Bastide et al., 2015; Cenci and Björklund, 2020; Chen et al., 2020). We employed it as a model of drug-induced long-term adaptations that manifest themselves at the behavioral level, with a prime interest in the underlying signaling mechanisms. These adaptations are by no means unique for LID and/or PD but are induced by other dopaminergic as well as non-dopaminergic drugs. Signaling plasticity could cause detrimental side effects limiting the therapeutic utility of drugs (e.g., in case of LID), or contribute to their addictive properties. A better understanding of the molecular mechanisms involved in these adaptations is necessary for the development of new or improvement of existing therapies. For example, the ability of the short Arr3-derived T1A peptide to substitute for the full-length Arr3 protein *in vivo* suggests the possibility of developing peptide-based tools with positive or dominant-negative action to control Arr3-dependent regulation of the JNK pathway for therapeutic purposes in human patients with a normal complement of WT Arr3.

## Acknowledgements.

We thank Dr. Robert J. Lefkowitz (Duke University) for the gift of the arrestin knockout mice. We are grateful to Dr. John F. Neumaier (University of Washington) for the gift of the rat enkephalin and dynorphin promoter vectors. This work was supported by NIH grants RO1 NS065868 and R21 DA030103 (to EVG) and R35 GM122491 (to VVG).

## MATERIALS AND METHODS

### Virus construction and preparation

The full-length coding sequence of the bovine Arr3 (gi 6978467) was N-terminally tagged with HA. The viral construct assembled in the lentiviral vector (LV) pLenti6.4/V5-DEST included GFP under control of the CMV promoter and downstream of it co-cistronic HA-tagged Arr3 under control of super-IRES (sIRES), which has been shown to significantly increase protein expression (Bochkov and Palmenberg, 2006). GFP alone was used as a control. The HA-tagged V343T mutant (Seo et al., 2011) and Venus-tagged T1A were produced by mutagenesis and verified by sequencing. The rat enkephalin (ENK) and dynorphin (DYN) promoter vectors were a gift from Dr. Neumaier (the University of Washington) (Ferguson et al., 2011). The promoter sequences were inserted into pLenti6.4/V5-DEST vector. The promoters were followed by sIRES before the sequence of HA-tagged Arr3. The lentiviruses were produced using the ViraPower system (Invitrogen, Carlsbad, CA), concentrated and purified as described (Ahmed et al., 2010). Viral titers of LVs with the CMV promoter were measured based on GFP or HA expression using HEK293 cells infected with the appropriate lentiviruses. The titers of lentiviruses with cell-specific promoters were determined by Western blot for HIV1 p24 band using mouse monoclonal antibody (ThermoFisher).

The adeno associated viruses serotype 5 (AAV5) encoding GFP (control) or HA-tagged Arr3 (with co-cistronic GFP) under control of the CMV promoter were constructed and produced using AAV-DJ Helper Free Expression System and 292AAV cell line (Cell Biolabs, Inc.). The viral tier was determined by standard qPCR with the BioRad SYBR Green supermix using serial dilutions of the cloning vector pAAV-MCS as standards.

### Mouse surgeries and virus injection

Adult wild type (WT), arrestin-2 knockout (A2KO), and Arr3 knockout (A3KO) mice bred on a C57Bl/6j background (Charles River) were used. A2KO and A3KO mice were a generous gift from Dr. R. J. Lefkowitz (Duke University). The animals were housed at the Vanderbilt University animal facility in a 12/12 light/dark cycle with free access to food and water. Mice were bred using heterozygous breeding pairs to obtain knockout and WT littermates. To maintain genetic homogeneity, mice were consistently backcrossed to WT C57Bl mice from Charles River. The same WT animals were used for backcrossing of A2KO and A3KO mice to minimize background genetic differences. All procedures were performed in accordance with the NIH *Guide for the Care and Use of Laboratory Animals* and approved by the Vanderbilt University IACUC.

The 6-OHDA lesion was performed essentially as described (Ahmed et al., 2007; Bychkov et al., 2007). Briefly, mice were deeply anesthetized with ketamine/xylazine (100/10 mg/kg i.p.) and mounted on a stereotaxis. Mice were treated with desimipramine (25 mg/kg i.p.) for 20 min prior to infusion of 6-hydroxydopamine (6-OHDA). 6-OHDA (2 μl; 2 μg/μl solution in 0.05% ascorbic acid) was infused unilaterally into the medial forebrain bundle at coordinates AP=1.1; ML=1.3; H DV=5.03. The mice were allowed to recover for 4 weeks after surgery.

The viruses were injected 24 h after the end of the behavioral pre-testing. The mice were anesthetized with ketamine/xylazine (100/10 mg/kg i.p.) and mounted on a stereotaxis. The virus injection (3 μl of the concentrated virus in saline per striatum 0.3 μl/min) was made unilaterally (on the 6-OHDA-lesioned side) into the dorso-lateral caudate-putamen at coordinates AP +1.02; ML 1.65; DV 3.55.

### Mouse behavior

#### Rotations

Four weeks after the 6-OHDA lesion, the animals were tested for rotational response to apomorphine (0.1 mg/kg s.c.) for 1 h using an automated rotometer (AccuScan Instruments). Both ipsilateral and contralateral 360° turns were recorded, and the net rotational asymmetry (contralateral minus ipsilateral turns) was calculated. The animals were then treated, starting the next day, with L-DOPA (L-3,4-Dyhydroxyphenylalanine methyl ester hydrochloride, 25 mg/kg i.p. twice daily, morning and afternoon) for 5 days, and the rotational response was recorded every day after the morning injection (pre-testing). Five days after the injection of lentiviruses or 3 weeks after the AAV injection, the mice were again treated with L-DOPA (25 mg/kg i.p. twice daily) and tested for L-DOPA-induced rotations every day for 10 days, as described (Ahmed et al., 2010; Ahmed et al., 2015a).

#### Abnormal Involuntary Movements (AIMs)

Testing for AIMs was performed essentially as described (Ahmed et al., 2010). The mice were treated with increasing doses of L-DOPA (2-10 mg/kg day) for 15 days, starting 4 weeks after the 6-OHDA lesion. The dose regiment was adjusted for each individual animal until substantial AIMs scores were achieved. The virus injection was performed on day 14, and the mice were allowed to recover for 5 days (all LV experiments) or 3 weeks (the LV-AAV experiment) before resuming the treatment for 18 more days with 10 mg/kg L-DOPA. AIMs were assessed by an independent observer in a blind manner on a 0-4 scale (Lundblad et al., 2002) on every third day, as described (Ahmed et al., 2010; Ahmed et al., 2015a). Briefly, the AIMs will be scored by an independent observer unaware of the animal’s experimental status on 0–4 scale: 0 – no AIMs; 1 – infrequent AIMs occurring < 50% of the time; 2 – frequent AIMs occurring >50% of the time; 3 – constantly present AIMs that are interrupted by external stimulation (tap on the cage); 4 – constant AIMs that are insensitive to external stimulation. Four types (orolingual, limb, locomotor, and axial) of AIMs will be assessed every 20 min for 1 min during a 3-h session every 3^rd^ day (total 9 observations; maximal score for each observation 16, maximal total score per session 144).

*The cylinder test* was performed as described (Ahmed et al., 2010). Half of the WT and A3KO mice were randomly selected to receive injections of L-DOPA (6 mg/kg); the other half received saline. The mice were placed in a glass cylinder close to a mirror and their behavior was video recorded under a red light for 6 min. The behavior was scored from videotapes by an independent observer blind to the mouse genotype, experimental condition, and the side of the lesion. The next day, the treatment groups were switched. The number of times a mouse used the contralateral (injured) or ipsilateral paw for support was counted from the recordings by an observer blind to the animal’s treatment (use is defined as the animal supporting its body with digits extended). Preferential use of the paw ipsilateral to the lesion (controlled by the intact hemisphere) is indicative of the akinetic defect of the lesion. Acute L-DOPA administration enhances the use of the contralateral paw, controlled by the lesioned hemisphere (Cenci and Lundblad, 2007; Lee et al., 2000; Mela et al., 2007). The results are expressed as a percentage of contralateral paw usage relative to the ipsilateral paw.

### Tissue preparation

Upon completion of the drug administration and behavioral testing, the mice were anesthesized with isoflurane, decapitated, the brains were collected and rapidly frozen on dry ice. The midbrain containing the substantia nigra was dissected and post-fixed in 4% paraformaldehyde for tyrosine hydroxylase (TH) immunohistochemistry to determine the loss of dopaminergic neurons. The rostral parts of the brains containing the striata were kept at −80°C until samples were collected for Western blot analysis. To collect the samples for Western blot, the brains were cut through the striatum, the position of the virus injection track was identified, and the tissue around the injection track was collected to determine the expression of virus-encoded proteins and TH level, as described previously (Ahmed et al., 2010; Ahmed et al., 2015a). Randomly selected animals were overdosed with ketamine/xylazine and transcardially perfused with 4% paraformaldehyde. The brains were removed, post-fixed in 4% paraformaldehyde, cryoprotected, kept frozen at −80°C, then cut into μM sections and used for immunohistochemistry.

### Western blot

To determine the lesion quality, we used Western blot of the striatal tissue for TH with rabbit anti-TH antibody (Chemicon, AB152; 1:10,000 dilution). The expression of GFP- or Venus-tagged constructs was measured by Western blot with mouse anti-GFP antibody (Clontech; 1:2,000). The expression of WT and mutant arrestin-3 was measured with rabbit polyclonal anti-arrestin-3 antibody, as described (Ahmed et al., 2007). To detect HA-tagged constructs, rabbit monoclonal anti-HA antibody (Cell Signaling, cat.# 3724; 1:1,000) was used. Active JNK was detected with rabbit antibodies to doubly phosphorylated (Thr183/Tyr185) (Cell Signaling, cat.# 4668 or 9251; 1: 1,000). c-Jun phosphorylated at the JNK phosphorylation site Ser73 was detected with phospho-specific rabbit antibody (Cell Signaling, cat.# 3270; 1:1,000) and the total c-Jun – with mouse monoclonal antibody (Cell Signaling, cat.# 2315; 1:1,000). Doubly phosphorylated (active) ERK1/2 was detected with a phospho-ERK-specific [phosphor-ERK1/2(Thr202/Tyr204)] mouse monoclonal antibody (Cell Signaling, cat.# 9106) at 1:2,000 dilution. After stripping, blots were re-probed with a rabbit anti-ERK1/2 antibody (Cell Signaling, cat.# 9102) at 1:1,000 dilution to detect total ERK1/2. Akt phosphorylated at Thr308 was visualized with rabbit anti-Akt(T308) antibody (Cell Signaling, cat.# 9275), phosphorylated at Ser473 - with anti-Akt(S473) rabbit antibody (Cell Signaling, cat.# 4060), and total Akt was detected with a rabbit antibody (Cell Signaling, cat.# 9272), all at 1:1,000 dilution. Phospho-p38 was detected with rabbit monoclonal antibody (Cell Signaling, cat.# 4511; 1:1,000) and total p38 – with rabbit antibody (Cell Signaling, cat.# 9212, 1:1,000).

Anti-rabbit or anti-mouse horseradish peroxidase-conjugated secondary antibodies (Jackson Immunoresearch) were used at 1:10,000 dilution. The blots were developed with a chemiluminescent substrate and either exposed to X-ray film or subjected to direct detection using C-DiGit Blot scanner (Li-Cor). The gray values of the bands on X-ray film were measured with a Versadoc imaging system (Bio-Rad) with QuantityOne software. Following direct detection Image Studio software was used.

Fluorescent detection of the peptide expression combined with TH and the marker of medium spiny neurons forkhead box P1 (FOXP1) (Tamura et al., 2004) was performed with mouse anti-GFP antibody (1:1,000), rabbit anti-TH (1:1,000) and rabbit anti-FOXP1 (1:500) followed by IRDye donkey anti-mouse 800CW (green) and donkey anti-rabbit 680 (red) antibodies (Li-Cor) at 1:10,000 dilution. The blots were imaged with the Odyssey Clx system and analyzed with Image Studio software.

### Immunohistochemistry

The lesion quality was tested by TH immunohistochemistry. The midbrain sections containing the substantia nigra were stained with rabbit anti-TH antibody (Chemicon, Temecula, CA; 1:1,000 overnight at 4°C) followed by anti-rabbit secondary antibody conjugated with Alexa594 (red).

To detect the expression of GFP-tagged constructs by immunohistochemistry, the mouse monoclonal anti-GFP (Jl-8, Clontech; 1:500 overnight at 4°C) primary antibody was used, followed by goat anti-rabbit biotinylated secondary antibody and streptavidin conjugated with Alexa Fluor 488 (Invitrogen, Carlsbad, CA). For double labeling with FOXP1, the sections were labeled with mouse anti-GFP (1:500) antibody and rabbit anti-FOXP1 antibody (Cell Signaling Technology; 1:400) and detected by biotinylated secondary antibody/streptavidin-Alexa Fluor 488 (GFP; green) and anti-rabbit-Alexa Fluor 568 (FOXP1; red). Alternatively, chicken anti-GFP antibody (Invitrogen; 1:500) were used. For double labeling with TH, the sections were labeled with mouse anti-GFP (1:500) antibody and rabbit anti-TH antibody (Chemicon; 1:1000) and detected by biotinylated secondary antibody/streptavidin-Alexa Fluor 488 (GFP; green) and anti-rabbit-Alexa Fluor 568 (TH; red). Double labeling for phospho-c-Jun and TH was performed with phospho-specific rabbit monoclonal antibody (Cell Signaling, cat.# 3270; 1:500) for c-Jun and sheep anti-TH antibody (Novex Biotechnology; 1:400), biotinylated secondary antibody/streptavidin-Alexa Fluor 488 (phospho-c-Jun; green) and anti-rabbit-Alexa Fluor 568 (TH; red).

The sections were photographed at low magnification on a Nikon TE2000-E automated microscope with a 4x dry objective and an Andor Zyla high resolution digital camera using the stitching function of the Nikon NIS-Elements software. High power photographs were collected on an Olympus FV-100 confocal microscope in the green and red channels with z-sectioning using a 40x oil immersion objective at 1024 x 1024 pixels. The images were assembled in Photoshop, with minimal contrast adjustments applied separately to channels to equalize their intensity.

### In-cell JNK phosphorylation assay

Human neuroblastoma SH-SY5Y cells were cultured in DMEM/F12 media supplemented with 10% fetal bovine serum and 1% penicillium/streptomycin. Cells were co-transfected with HA-tagged JNK3α2 and with either: (1) empty vector; (2) WT arrestin-3; or (3) arrestin-3-V343T mutant. Similarly, in the T1A experiment the cells were transfected with HA-JNK3α2 and with half or full concentration of WT Venus-tagged-ARR3, half or full concentration of Ve-T1A, or full concentration of Ve-B1A. In the arrestin-3 dose-response experiment, the cells were transfected with increasing concentrations of WT arrestin-3. Activation of JNK3 was assessed by Western blot with a ppJNK specific antibody (Cell Signaling, cat. # 9251 or 4668), as described (Chen et al., 2017; Zhan et al., 2015; Zhan et al., 2016).To detect the JNK3 expression, JNK3-specific mouse (Santa Cruz Biotechnology) or rabbit anti-HA antibody (Cell Signaling, cat. # 3724) was used. Arr3 and Arr3-derived constructs were detected with rabbit anti-Arr3 (Ahmed et al., 2007) or mouse anti-GFP (Clontech JL-8) antibody. Total JNK level was measured with rabbit anti-SAPK/JNK antibody (Cell Signaling technology, cat.# 9152).

### Immunoprecipitation from the striatal tissue

The lesioned and intact striata from 6-OHDA-lesioned WT and A3KO mice chronically treated with L-DOPA and challenged with a L-DOPA dose 45 min before sacrifice were dissected and lysed in NP-40-containing immunoprecipitation (IP) buffer, as described (Ahmed et al., 2015b; Ahmed et al., 2011). The lysates were cleared by centrifugation, equalized by protein content with IP buffer, and the supernatants were incubated overnight at 4°C with mouse anti-JNK3 antibody (Santa Cruz Biotechnology). Protein G slurry (20 μl) was added to each tube, and incubation was carried out at 4°C for 2 h. The resin was washed 3 times with IP buffer, and samples were eluted with SDS buffer. Samples were analyzed by Western blot with rabbit anti-ppJNK (Cell Signaling Technology, cat. #4668) and rabbit anti-JNK3 (Cell Signaling Technology, cat. #2305) antibodies.

### Data Analysis

The rotation data were analyzed by two-way repeated measure ANOVA with Group (GFP versus Arr3 and Arr3 mutant, or T1A or B1A peptide) as a between group and Day as a repeated measure factor. The group differences across sessions were assessed by the post hoc Bonferroni/Dunn test with correction for multiple comparisons or by Dunnett’s test where appropriate. When a significant effect of Group was observed, the data for individual sessions were compared by the unpaired Student’s test (two groups) or by the Bonferroni/Dunn post hoc test, when appropriate. The AIMs scores for each session were compared by Kruskal-Wallis non-parametric ANOVA followed by Dunn’s post hoc test with correction for multiple comparisons. The data for the cylinder test were analyzed using the nonparametric paired Wilcoxon Signed Rank test (to evaluate the effect of L-DOPA) or the Mann-Whitney test (for group comparison). The Western blot data were analyzed by ANOVA with Genotype, expressed protein, and/or Treatment as main factors, with Hemisphere treated as a within group factor, where appropriate. StatView (SAS Institute) and Prizm GraphPad software were used for the statistical analysis. The value of p<0.05 was considered significant.

