## Supplementary material for "Arrestin-3-assisted activation of JNK3 mediates dopaminergic behavioral and signaling plasticity in vivo": Supplemental Information - Ahmed et al..docx

**SUPPLEMENTAL FIGURES**

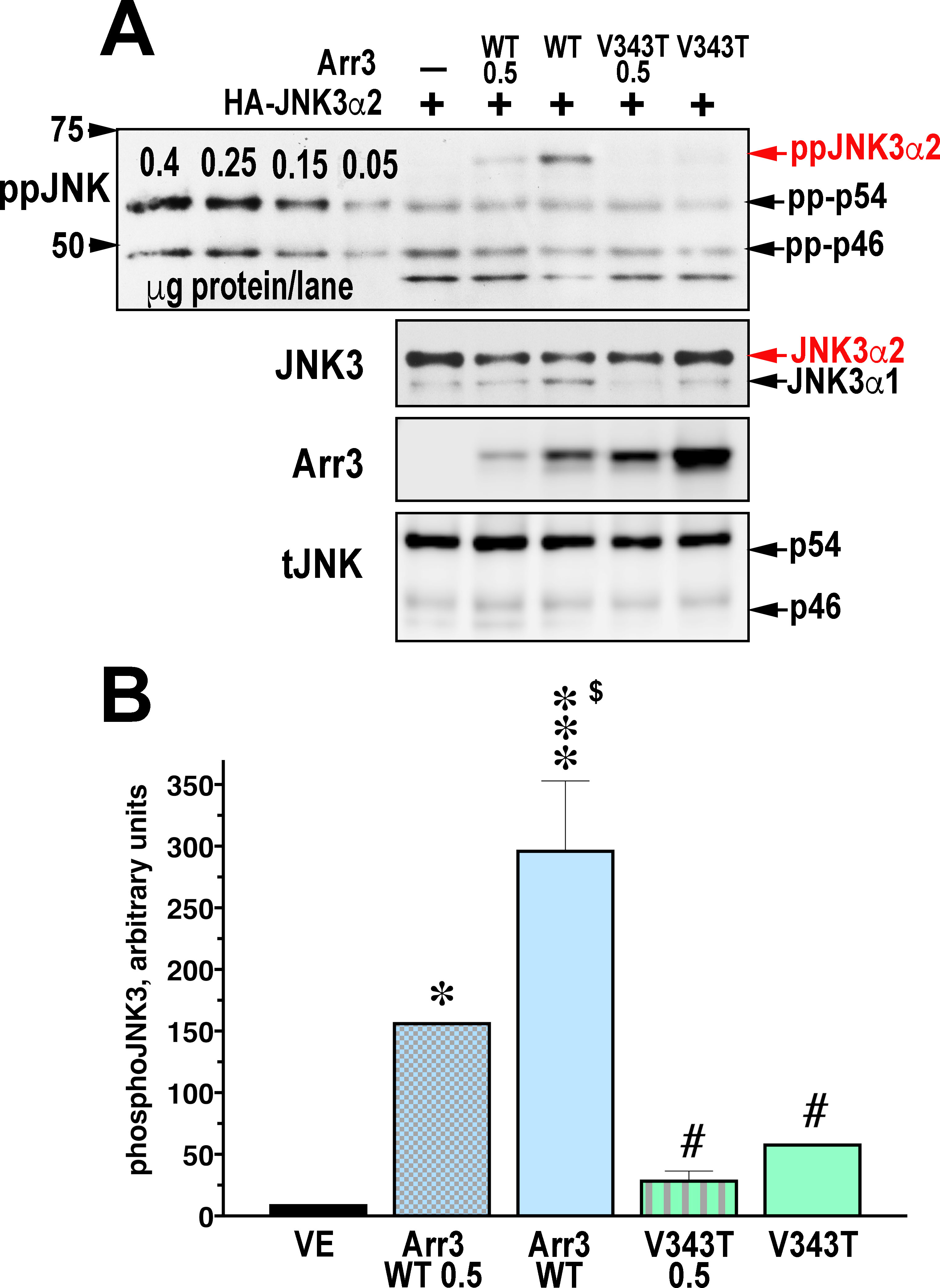

**Figure S1. Arr3-V343T mutant has reduced ability to activate JNK3 in neuronal cells.** (**A**) Human neuroblastoma SH-SY5Y cells were co-transfected with HA-JNK3α2, and with two concentrations of either WT Arr3 or the Arr3-V434T mutant (0.5 - half the amount of DNA used for transfection). Representative Western blots show the level of JNK phosphorylation and the expression of transfected JNK3α2 as well as endogenous JNK3α1 visualized with anti-JNK3 antibody. Arr3 and Arr3-V343T were detected with rabbit polyclonal anti-Arr3 antibody (*6*). The total level of JNK is shown as loading control. Left 4 lanes - serial dilutions of the lysate of anisomycin-stimulated HEK293 cells. **(B)** Quantification of the level of JNK3α2 phosphorylation from 5 independent experiments. WT Arr3 significantly increased JNK3 phosphorylation, as compared to control, whereas Arr3-V434T had minimal effect. *** - p<0.01, * - p<0.05 to VE; $ - p<0.05 to Arr3 0.5; # - p<0.01 to Arr3 by Bonferroni post hoc comparison with correction for multiple comparisons following one-way ANOVA with Protein as main factor (p<0.0001).

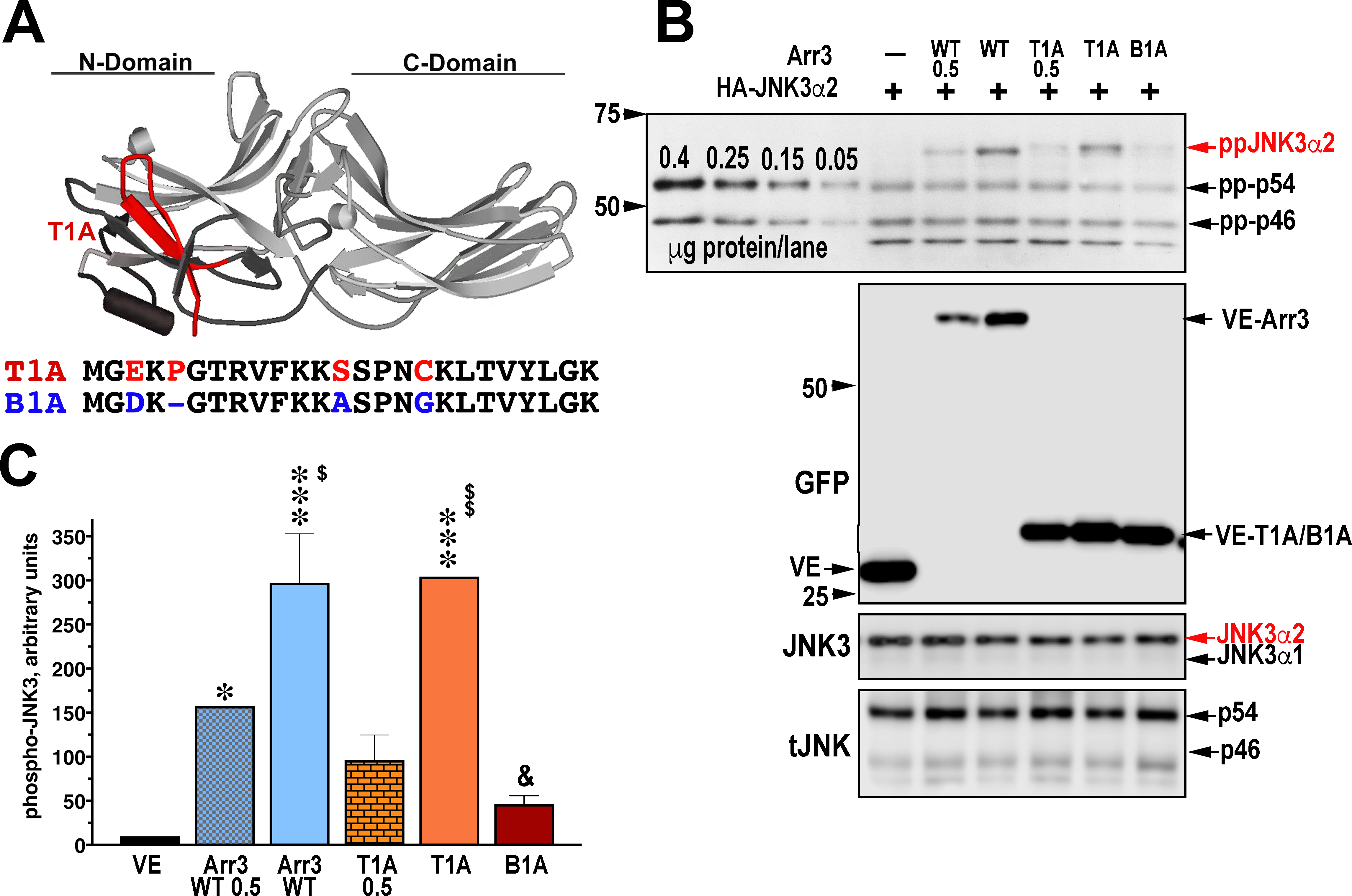

**Figure S2. (A**) Structure of Arr3 with T1A peptide highlighted in red. Below are the amino acid sequences of T1A and of the corresponding peptide B1A derived from Arr2 with mismatches highlighted in red (in T1A) and blue (in B1A). (**B**) Representative Western blots show JNK3 activation by the full length Arr3 (WT) and T1A (both at full and half concentrations, (0.5)) and minimal activation by B1A at full concentration. Also shown are the expressions of transfected proteins, and the total JNK as loading control. Left four lanes in the phosphor-JNK blot show serial dilutions of the lysate of anisomycin-stimulated HEK293 cells as positive control. (**B**) Quantification of the ppJNK3α2 level from four independent experiments. *** - p<0.001, * - p<0.05 to VE; $$ - p<0.01, $ - p<0.05 to corresponding half expression of Arr3/T1A; & - p<0.01 to full expression of T1A and Arr3 by Bonferroni post hoc comparison following one-way ANOVA with Protein as main factor (p<0.0001).

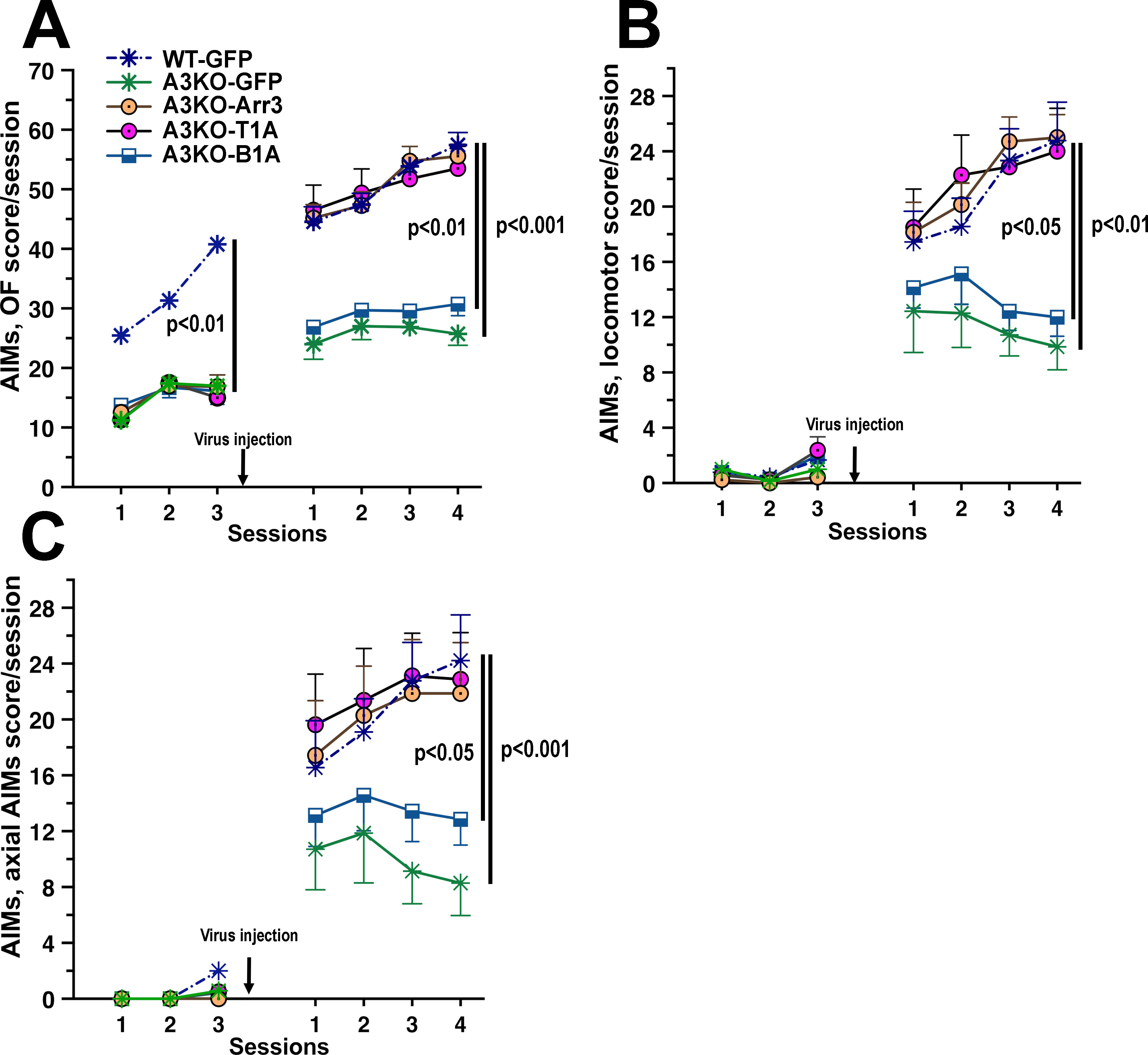

**Figure S3. T1A and full-length Arr3 fully rescue all categories of AIMs in A3KO mice**. AIMs are classified into four categories, orolingual, limb, locomotor, and axial, which are scored independently. **(A)** Combined score for orolingual and limb AIMs. **(B)** Scores for locomotor AIMs. **(C)** Scores for axial AIMs. Arrow shows the time of the virus injection. Significance shown applies to the last testing session and is for the comparisons between each of the three groups (WT, A3KO-Arr3, A3KO-T1A) and either A3KO-GFP or A3KO-B1A by Dunn’s post hoc test following Kruskal-Wallis non-parametric ANOVA. Note early appearance of OF AIMs as compared to the locomotor and axial AIMs and the tendency to a reduction in frequencies across post-injection sessions of both locomotor and axial AIMS in A3KO-GFP mice.

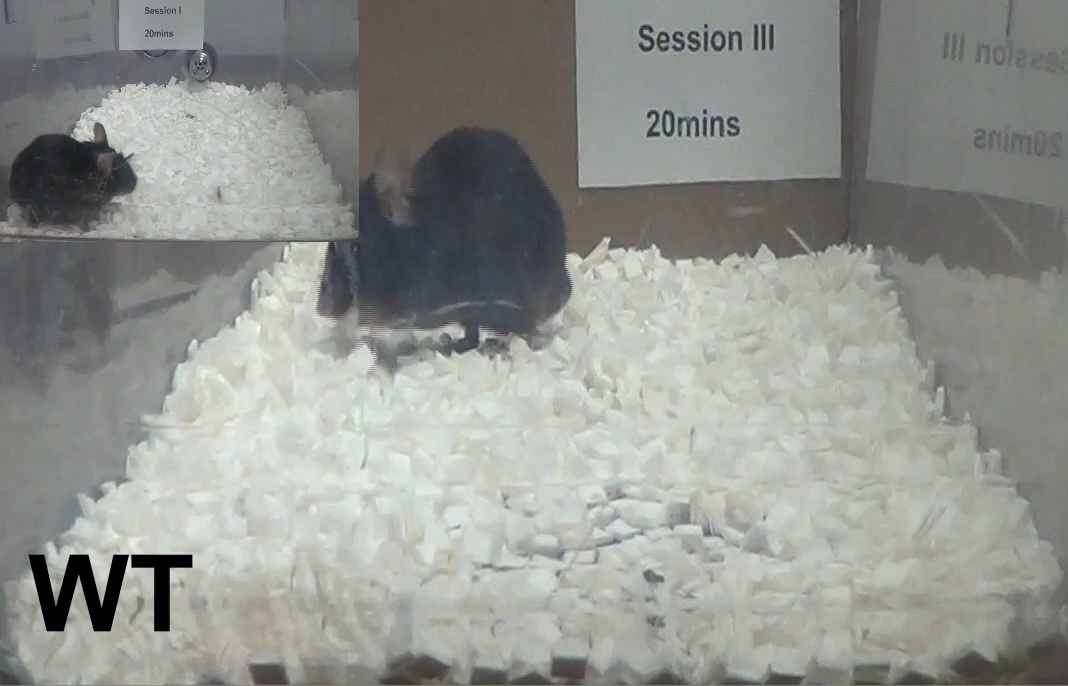
**Video S1. Representative AIM sessions in a wild type control (GFP) mouse.** Shows sensitization of AIMs behavior a wild type mouse from the first L-DOPA treatment in Session I (insert on the left; the mouse behaves normally) to intense orofacial, limb, and locomotor AIMs in post-injection Session III.

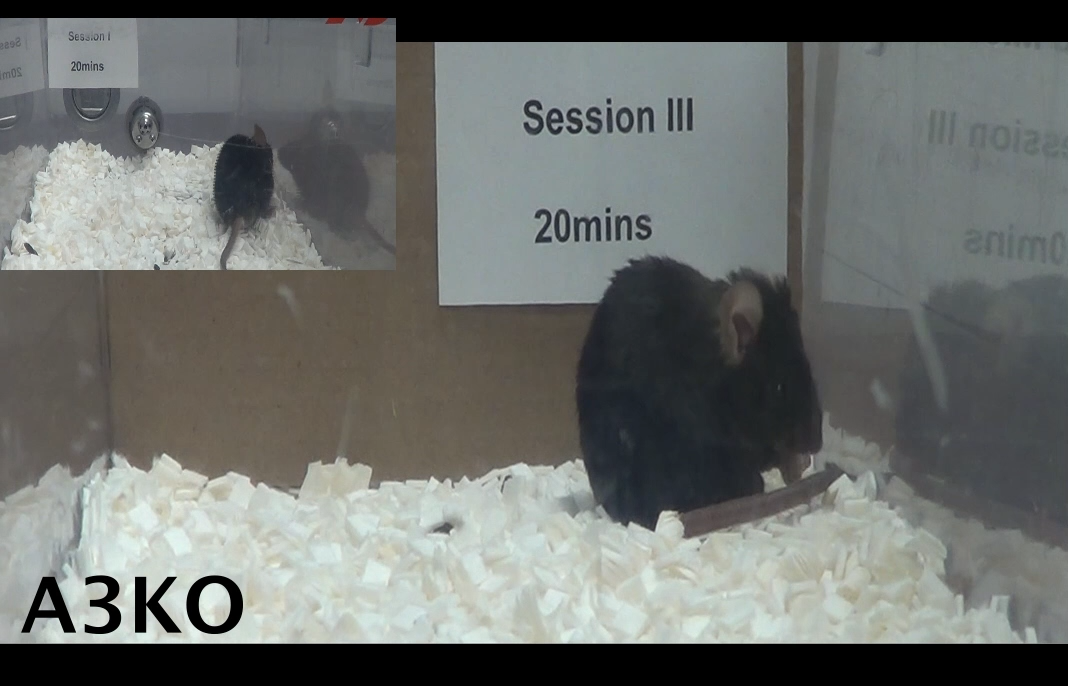

**Video S2. Representative AIM sessions in a A3KO control (GFP) mouse.** Show reduced sensitization of AIMs behavior in a A3KO mouse. In post-injection Session III, the mouse shows only orofacial and limb, with minimal locomotor AIMs. Insert on the left – pre-injection Session I.

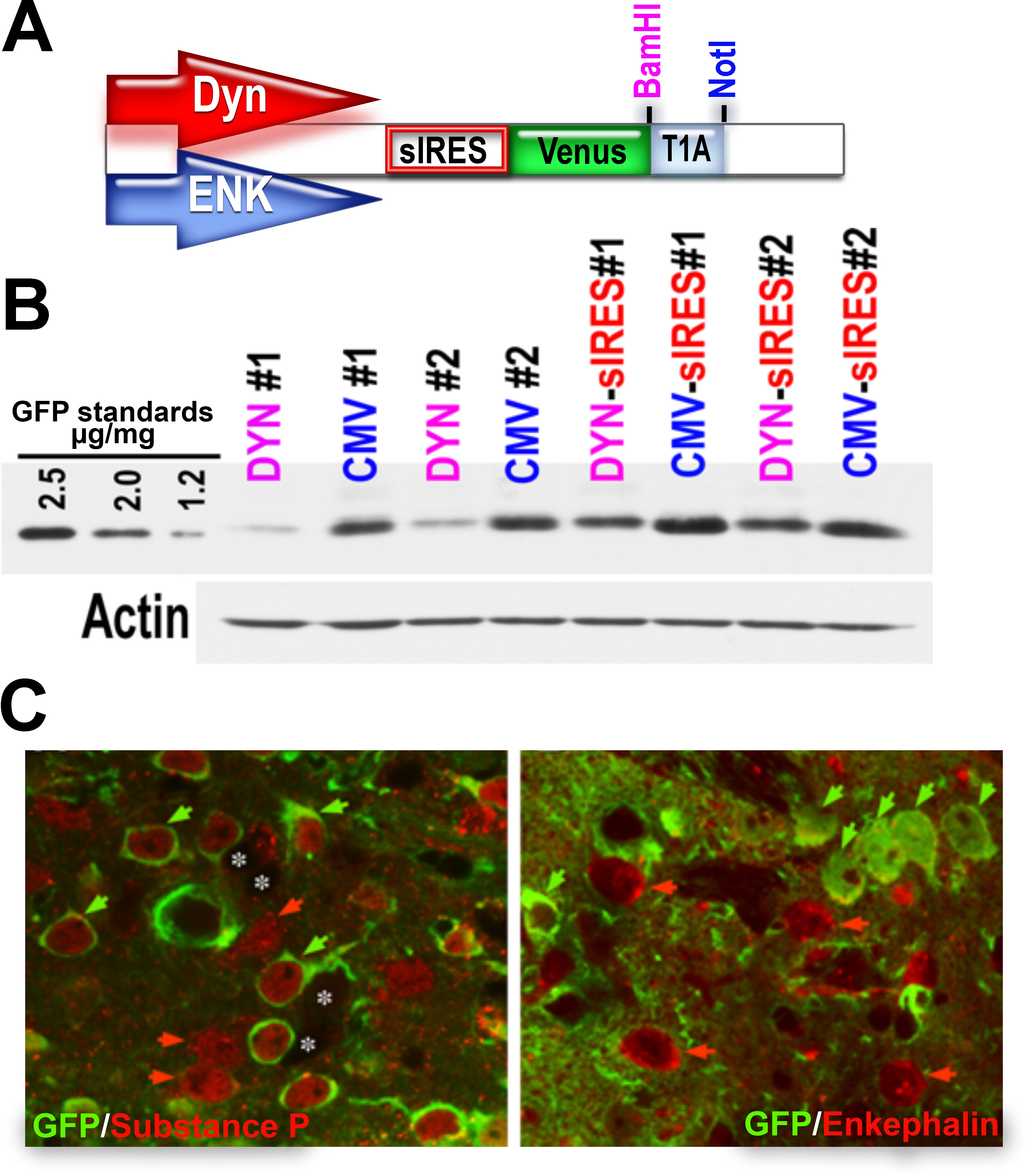

**Figure S4. Expression of T1A in the striatum driven by neuron-specific promoters.** **(A)** Schematic of the structure of the lentiviruses used to express the Arr3-derived T1A peptide in a neuron-specific manner. To increase the expression levels and to equalize the expression driven by different promoters, the constructs were subcloned under the control of superIRES (sIRES). **(B)** Direct side-by-side comparison of the level of transgene expression in the striatum achieved by lentiviral transfer under the control of the CMV and dynorphin (DYN) promoters with or without sIRES. Mice were injected with lentiviruses encoding Venus-T1A under the control of either the CMV or DYN promoters with or without sIRES and sacrificed 5 days later. The striatal samples were collected, equalized by total protein, and processed for Western blot with anti-GFP antibodies. Note the considerably lower expression under the DYN promoter alone as compared to CMV alone and the comparable levels achieved when DYN-sIRES and CMV constructs were used (DYN only drives expression in approximately 50% of infected striatal neurons). **(C)** Mice were injected with lenti-DYN-Venus-T1A into the CPu and transcardially perfused 7 days later; the brains were postfixed, cryoprotected, frozen, and sectioned on a cryostat at 30 μm. Free-floating sections were stained with mouse anti-GFP antibody (Clontech) (green) and rabbit antibody to substance P (left image) or enkephalin (right image) (red). Left image: Green arrows point to substance P-positive neurons expressing Venus-T1A; red arrows - to substance P-positive uninfected neurons, and white asterisks indicate substance P-negative neurons that do not express Venus-T1A. Right image: Green arrows point to enkephalin-negative neurons expressing Venus-T1A; red arrows - to enkephalin-positive neurons that do not express Venus-T1A.

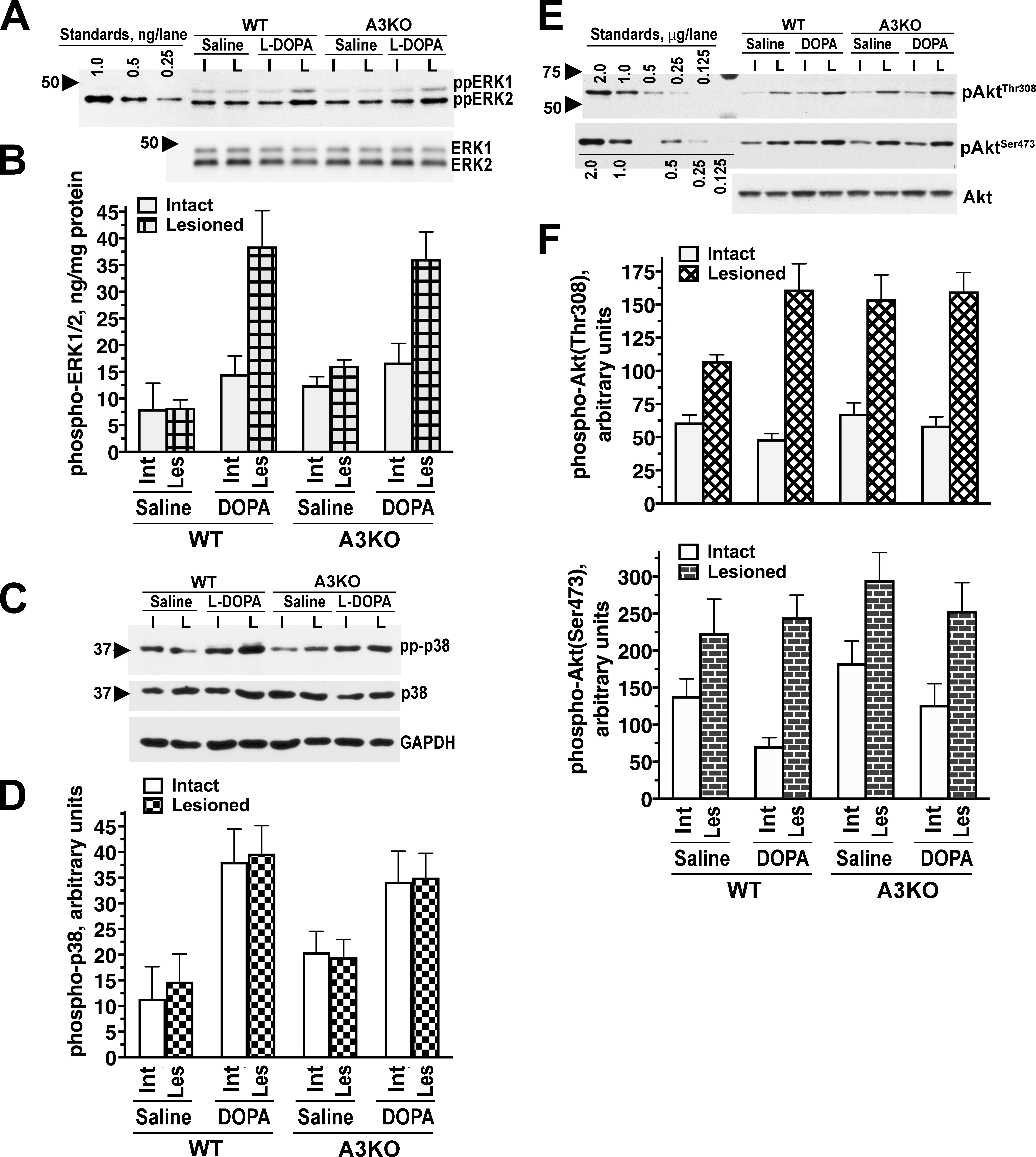

**Figure S5. Signaling in the lesioned striata in WT and A3KO mice.** The WT and A3KO mice were treated with L-DOPA for 10 days and then challenged with either saline or L-DOPA 45 min before sacrifice as described in Methods. The data were analyzed by three-way repeated measure ANOVA with Genotype and Challenge as between group and Hemisphere as within group factors. **(A)** Representative Western blot showing the level of ERK1/2 activation in the lesioned striata of WT and A3KO mice chronically treated with L-DOPA and then challenged with saline or L-DOPA 45 min prior to sacrifice. Left 3 lanes - different concentrations of the purified human ppERK2 used as standards. Note the significantly higher level of ERK1/2 phosphorylation (ppERK1 and ppERK2) in the lesioned striatum in response to the drug challenge but not to saline, with the total ERK1/2 level remaining the same**. (B)** Quantification of the Western blot data (N=10-12) demonstrating a comparable degree of ERK super-activity in the lesioned striatum of WT and A3KO in response to L-DOPA challenge. For pp-ERK1/2, the main effect of Genotype was not significant (p=0.56) and neither was the Genotype x Hemisphere interaction (p=0.72) or Genotype x Challenge (p=0.55), indicating a similar degree of super-sensitivity to L-DOPA challenge in the lesioned striatum. **(C)** Representative Western blot showing the level of p38 activation in the lesioned striata of WT and A3KO mice. **(D)** Quantification of the Western blot data (N=10-12) demonstrating a comparable degree of p38 super-activity in WT and A3KO mice in response to L-DOPA challenge. For p-p38, the main effect of Genotype was not significant (p=0.56) and neither was the Genotype x Hemisphere (p=0.72) or Genotype x Challenge (p=0.35) interaction, indicating a similar degree of super-sensitivity in the intact and lesioned striatum in both WT and A3KO mice to L-DOPA challenge. (**E**) Akt signaling in the lesioned striata in WT and A3KO mice. Representative Western blot showing the level of Akt activation in the lesioned striata of WT and A3KO mice. Left 5 lanes - serial dilutions of the lysate of FGF-treated HEK293 cells in μg of total protein/lane. Note the significantly higher level of Akt phosphorylation at the main activating residue T308 (upper panel) as well as at the supplemental residue S473 (middle panel) regardless of the challenge, with the total Akt level remaining the same. **(F)** Quantification of the Western blot data (N=8-11) demonstrating a comparable degree of Akt super-activity in the lesioned striatum of WT and A3KO mice. The upper graph shows Akt phosphorylation at Thr308, the lower graph - at Ser473. The effect of Genotype was not significant for either T308 (p=0.27) or Ser473 (p=0.22). The Genotype x Hemisphere (p=0.72) or Genotype x Challenge (p=0.35) interactions were also not significant (p>0.05). For both Akt(Thr308) and Akt(Ser473), the only significant effect was that of Hemisphere (p<0.0001 for both), indicating increased Akt activation in the lesioned as compared to the intact striatum regardless of the challenge (Challenge main effect p=0.27 and p=0.43, respectively). Thus, there was no difference between WT and A3KO mice in the super-responsiveness of Akt in the lesioned hemisphere.

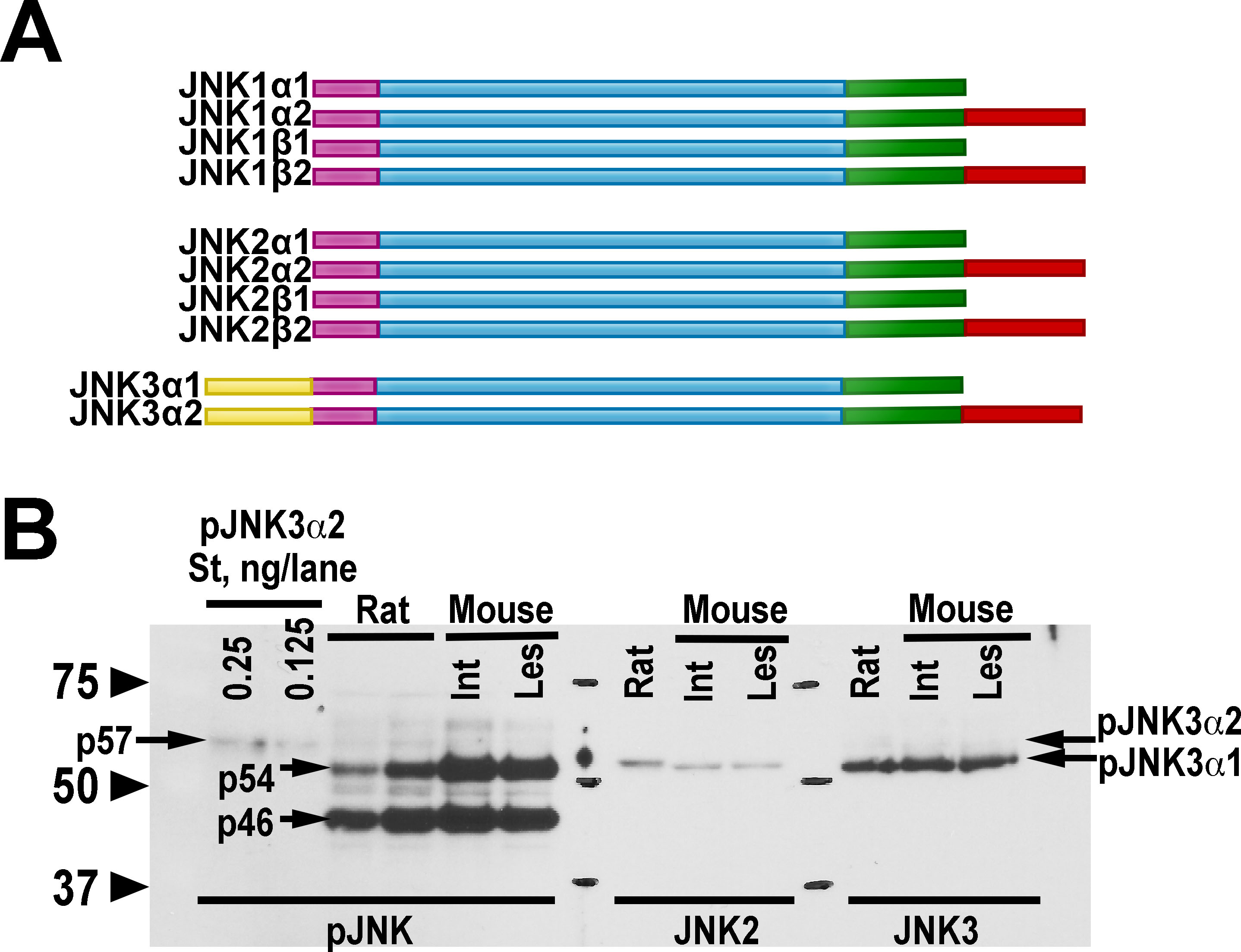

**Figure S6. The JNK isoforms in the striatum.** **(A)** Known isoforms/splicing variants of JNK1, JNK2, and JNK3 proteins. All 3 JNK genes (JNK1, JNK2, and JNK3) encode multiple splice variants. There are two variable regions in JNK1/2 and three in JNK3. All these alternative splicing products combine freely and yield four isoforms for JNK1 and JNK2. For JNK3, there are 2 known isoforms: the longer JNK3α2 and shorter JNK3α1. JNK3α2 is the longest isoform (57 kD) due to the N-terminal as well as C-terminal extensions. Red, C-terminal flexible extension; yellow, N-terminal flexible extension. **(B)** Expression of the JNK isoforms in the rodent striatum. The samples were probed for JNK activation with antibody detecting double phosphorylated JNK (left two lanes – purified phospho-JNK3α2 used to indicate the position of this isoform). Note two characteristic bands, p54 and p46. Also note that the longer p57 JNK3α2 isoform is barely detectable in the rat whole striatal lysate (two left lanes, 1x and 2x loads) and not detectable in the mouse lysate. Next two lanes show the samples probed with antibody specific for JNK2 longer isoforms (JNK2α2/JNK2β2), and the last two lanes – for JNK3. The shorter p54 JNK3α1 is the major JNK3 isoform in the rat and the mouse striatum.

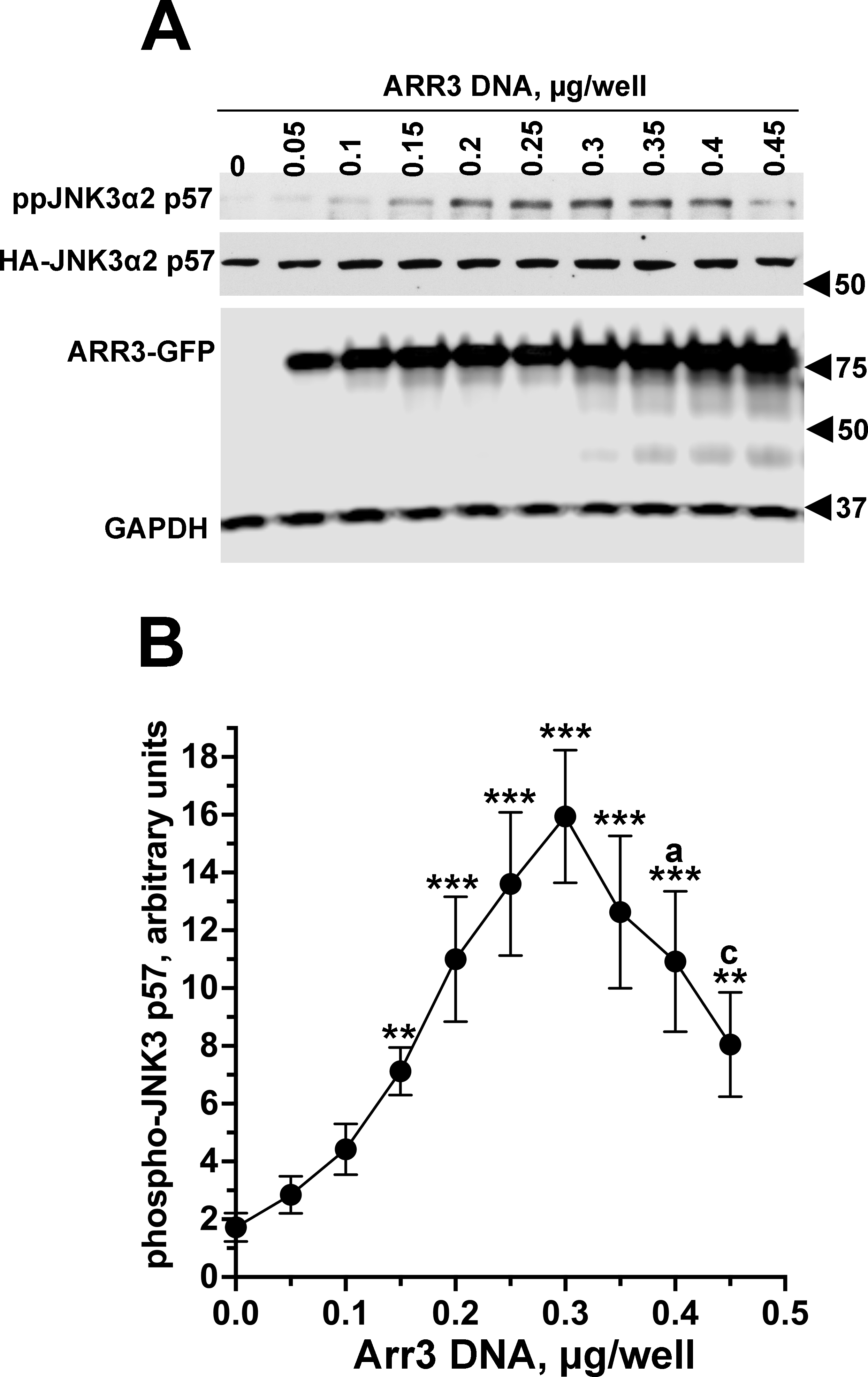

**Figure S7. Scaffolding action of Arr3 for the JNK activation cascade in neuronal SH-SY5Y cells. In-cell JNK3 phosphorylation in the presence of increasing concentrations of GFP-tagged Arr3.** SH-SY5Y cells were co-transfected with HA-tagged JNK3 and increasing amounts of GFP-tagged Arr3. **(A)** Cells were lysed 48 h post-transfection and lysates were blotted for phospho-JNK, HA (to detect HA-tagged JNK3α2 p57), GFP, and GAPDH. **(B)** Quantification of Western blot data for phosphor-JNK3α2 p57 band. ** - p<0.01, *** - p<0.001 to 0 point; a – p<0.05, c – p<0.001 to 0.3 (maximum) point by Tukey’s multiple comparison test following one-way ANOVA with Arr3 DNA amount treated as a factor.
